# Evolution of structural dynamics in bilobed proteins

**DOI:** 10.1101/2020.10.19.344861

**Authors:** Giorgos Gouridis, Yusran A. Muthahari, Marijn de Boer, Konstantinos Tassis, Alexandra Tsirigotaki, Niels Zijlstra, Nikolaos Eleftheriadis, Ruixue Xu, Martin Zacharias, Douglas A. Griffith, Yovin Sugijo, Alexander Dömling, Spiridoula Karamanou, Anastasios Economou, Thorben Cordes

## Abstract

Novel biophysical tools allow the structural dynamics of proteins, and the regulation of such dynamics by binding partners, to be explored in unprecedented detail. Although this has provided critical insights into protein function, the means by which structural dynamics direct protein evolution remains poorly understood. Here, we investigated how proteins with a bilobed structure, composed of two related domains from the type-II periplasmic binding protein domain family, have undergone divergent evolution leading to modification of their structural dynamics and function. We performed a structural analysis of ~600 bilobed proteins with a common primordial structural core, which we complemented with biophysical studies to explore the structural dynamics of selected examples by single-molecule Förster resonance energy transfer and Hydrogen-Deuterium exchange mass spectrometry. We show that evolutionary modifications of the structural core, largely at its termini, enables distinct structural dynamics, allowing the diversification of these proteins into transcription factors, enzymes, and extra-cytoplasmic transport-related proteins. Structural embellishments of the core created new interdomain interactions that stabilized structural states, reshaping the active site geometry, and ultimately, altered substrate specificity. Our findings reveal an as yet unrecognized mechanism for the emergence of functional promiscuity during long periods of protein evolution and are applicable to a large number of domain architectures.

## Introduction

Proteins drive and maintain all fundamental cellular processes^1^ by interactions with small molecules and/or other biopolymers. Important aspects of mechanistic information on proteins is accessible via structural analysis of their functional cycle.^2^ While classical approaches rely on the interpretation of static structure snapshots, the visualization of structural dynamics, i.e., to follow the interconversion of distinct structural states at high spatial and temporal resolution^3–7^, has been recognized as an essential complement. Emerging from this is the folding funnel model^8^ rooted in the free energy landscape theory^9–11^ that by now became widely accepted as a meaningful way to describe the ensemble of such states.^12–14^

Distinct structural states can originate from local flexibility, i.e., bond vibrations, side-chain rotations, loop motions (Fig. 1A, Tier-2 dynamics), changes in secondary structure (Fig. 1A, Tier-1 dynamics) or large-scale domain motions (Fig.1A, Tier-0 dynamics). The free energy landscape of a protein defines the lifetime of the structural states, ranging from nanoseconds (local flexibility) to seconds (large scale motions). Transitions between the states are referred to as conformational changes and are induced by interactions with ligands, post-translational modifications (e.g., phosphorylation) or chemical events such as nucleotide hydrolysis. The coupling of the latter to structural changes enables proteins to perform a diverse range of functions.

**Figure 1.**
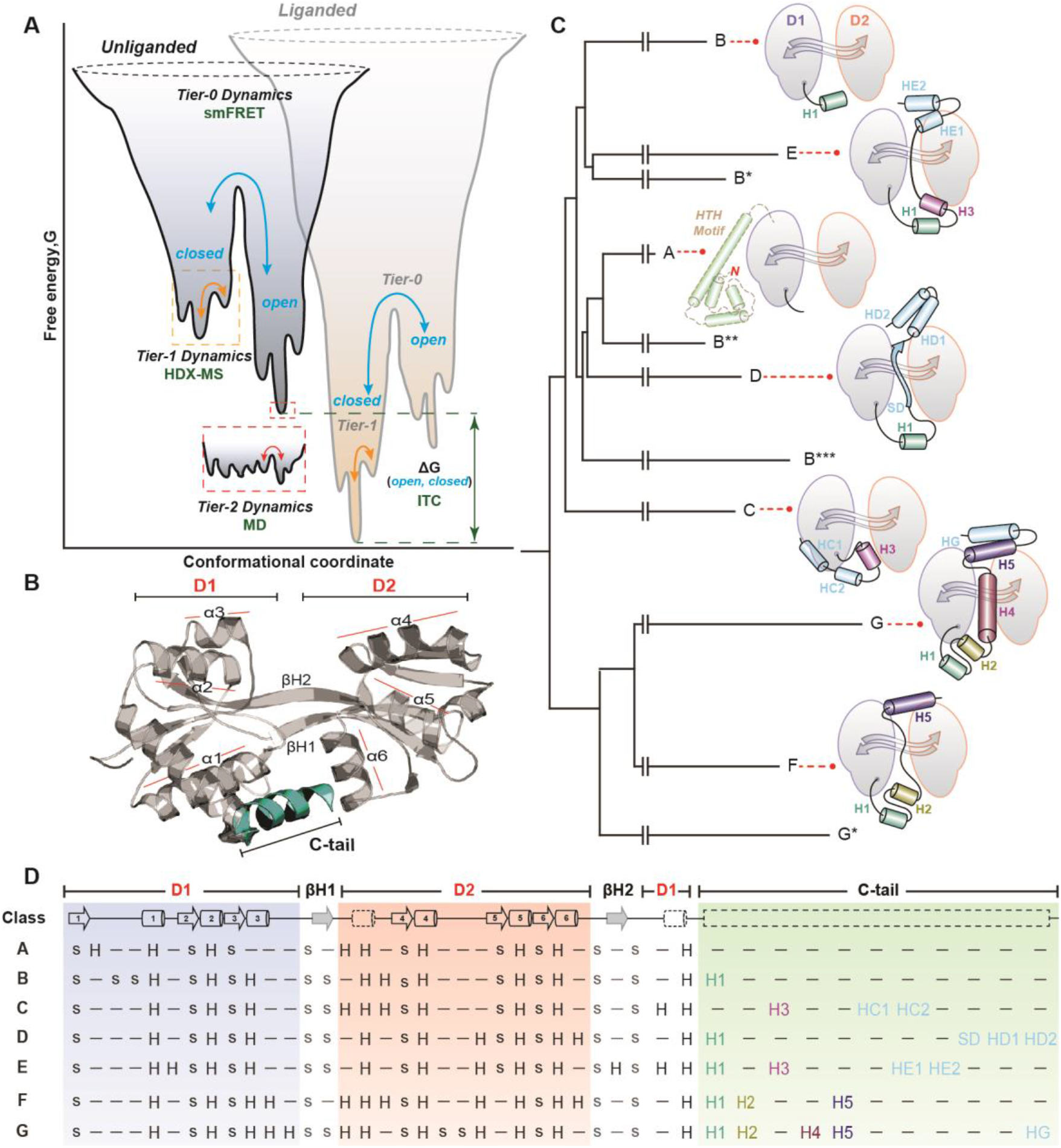
Energetic funnel, structure and evolution of the *cherry-core*. **(A)** One-dimensional cross section of a hypothetical protein energy landscape adapted from Kern and co-workers^12^ according to the Tier description and definitions introduced by Frauenfelder and co-workers.^15^ A structural state is defined as the lowest point of a well on the energy surface. The populations of the Tier-0 states “closed” and “open” are defined as Boltzmann distributions and their relative probabilities (p_C_, p_O_) are determined in this paper by smFRET, which allow to follow large-scale domain motions. Tier-1 states describe local and fast structural fluctuations (e.g., changes in secondary structure elements like loop motions or loss of secondary structure). Tier-1 dynamics were probed in this study by HDX-MS. Rapid and localized Tier-2 dynamics (e.g., side chain rotations) were not considered here, but can be monitored via MD simulations. Changes in the chemical environment, i.e., absence or presence of a ligand, modify the energy landscape via a bias for one of the two states. Typically, thermodynamic parameters such as the free energy difference of Tier-0 wells (ΔG_OC_) can be determined by ITC. **(B)** Structure of a representative bilobed protein, the substrate binding domain 2 (SBD2) of the ATP-binding cassette (ABC) amino acid transporter GlnPQ from *Lactococcus lactis* (PDB: 4KR5). **(C)** Summary of the structure-based phylogenetic tree with schematic representations of the different structural classes (class A-G) highlighting their termini. Complete sequence and structure-based phylogenetic trees are provided in Fig. S1 and SuppFile. 1, 2. We used the following notation: Secondary structure elements that are common between the different classes were assigned a number identifier (e.g. helix H5 in classes F and G), whereas the unique elements have a letter followed by a number identifier (e.g. HD1, first unique helix of class D proteins). An asterisk (*) marks a structural subclass (shown in detail in SuppFile. 3). Schematics are based on the crystal structures of the selected proteins. **(D)** Topology of bilobed proteins depicting the consensus of secondary structure elements: strands (s) or helices (H) belonging to the Domains (D1, D2), β-strands (β1, β2) and C-tails of all seven classes (A-G) are shown. Revised alignments of bilobed proteins are shown in SuppFile. 3. The secondary structure elements forming the conserved *cherry-core* structure are depicted on the top row.

Tier-0 dynamics were observed and characterized in various settings, e.g., in motor proteins, where they are used in propelling movement along filaments^16^, in the transport of molecules or biopolymers across biological membranes^17–21^, or in the activity of proteins that perform mechanical work.^22^ Tier-1 dynamics drive the actions of various signaling proteins for transmission of signals.^23–25^ Structural and biochemical data indicate that also enzymes show varying degrees of structural dynamics^26^, although it is not well understood what precise role this plays for catalytic activity. The current belief is that extensive structural dynamics in enzymes are not necessarily required for catalysis^27^, but rather enact in regulation. For instance, many protein kinases exploit Tier-0 dynamics to generate active or inactive structural states.^28^ Tier-2 dynamics have been shown to be important for the evolution of enzymatic function.^29^ In addition to domain motions occurring within a structure, protein oligomerization and the possible quaternary dynamics can also be relevant for function.^30^ A well-characterized example is the allostery of haemoglobin that occurs on the transition between two distinct quaternary states (relaxed and tense).^31,32^ All this highlights that multi-Tier dynamics of proteins occur on various time- and length-scales (Fig. 1A) and are often the basis for function. It is, however, not well understood how structural dynamics are designed and optimized during evolution to tailor protein function.

Analysis of protein sequences and structures has provided important insights into the evolution of protein function.^33^ Proteins can be decomposed into common constituent domains, a domain being a contiguous polypeptide chain within a protein that can fold and function independently. A powerful approach is to assign the domain components of proteins to families and superfamilies on the basis of sequence alone (Pfam database)^34^ or in combination with structural information (CATH and SCOP databases).^35,36^ The CATH and SCOP databases combined have identified ~3000 domain superfamilies comprising >50 million domains that account for ~70% of domains in completed genomes.^37^ More recently, the database ECOD groups domains by considering their evolutionary relationships.^38^ In ECOD ~760000 domains have been assigned to 3700 homologous groups.^39^ The most highly populated domain superfamilies/homologous groups are universal to all kingdoms of life.^40,41^ The prevalence of proteins with two or more domains, i.e., proteins with multidomain architectures, and the recurrent appearance of the same domain in non-homologous proteins suggests that functional domains are reused when creating new proteins and functions.^42^ Analyses of selected domain groups^43,44^, and more recent large-scale investigations^45,46^ have shown that domains in such groups generally share a common structural core (40-50% of the domain) that is highly conserved, even for relatives separated by billions of years. To achieve functional promiscuity of such a structural core, it has been proposed that larger structural embellishments (secondary structure elements or even entire domains) need to be added to the core for altered biochemical function.^47^ Such fusions occur frequently at the N or C termini^40^ which has led to the belief that structural elements act as Lego bricks, which are recombined in various ways for new functions to emerge during evolution.^42,47^

Much less is known about the evolution of structural dynamics or the role played by structural dynamics in the evolution of protein function. A recent model suggests that the native state of an evolved protein is the most abundant state of all possible structural states, which was selected for a specific function.^48,49^ The “avant-garde” evolvability theory proposes the existence of a highly promiscuous primordial protein structure. It is assumed that for the emergence of the evolved native state, several synergistic rounds of neutral mutations and plastic modifications were required, during which, the ability to evolve (evolvability) was traded for ligand-functional specificity.^50,51^ These observations were experimentally verified^52^ and agree with the proposal that changes in structural dynamics serve as a mechanism for the evolution of specialist enzymes from promiscuous generalists.^53^ Though, discordant examples are also observed in evolution.^54^ Neutral mutations do not compromise the native state and the original function of a protein, whereas plastic modifications are referred to as adaptive structural changes in response to a few mutations leading to functional specialization. Other recent studies indicate that this evolvability theory can explain the short-period evolution, e.g., variation of enzyme local flexibility (variation of Tier-1/2 dynamics) to acquire new functions, particularly well.^48–50,55^ Remarkably, this short-period evolution can be faithfully reproduced *in vitro* using directed evolutionary approaches based on consecutive rounds of single point mutations.^48,55^ Ancestral protein reconstruction has also been useful in elucidating the role of structural dynamics in the emergence of specialized amino acid binding proteins from a promiscuous ancestor.^56^ However, it still remains unclear how during longer periods of evolution, a primordial core structure evolves to modulate or diverge Tier-0 or quaternary dynamics and with that “generates” completely new functionalities.

Here, we test this aspect of the evolvability theory for proteins separated by long evolutionary periods during which Tier-0 or quaternary dynamics were introduced to an existing protein core structure. For this, we identified proteins with a bilobed domain structure with a high degree of plasticity and neutrality^29,49^, having (i) a conserved core-structure, (ii) large sequence diversity and related functional divergence and (iii) occurrence in all kingdoms of life. The selected structural core is composed of two Rossman folds, believed to be amongst the most ancient architectures^57^, connected by a single β-sheet (Fig. 1B). We analysed ~600 proteins with this structural core that contain two related domains from the type-II periplasmic binding protein (PBP) domain homologues (ECOD: X-, H- and T-groups Periplasmic Binding Protein-like II). We show here that different members of these homologues domains have undergone divergent evolution by the acquisition of domains or secondary structure elements, predominantly at their termini (Fig. 1C, D). Using a combination of structural analysis and biophysical investigations, we demonstrate that such structural embellishments confer multi-Tier conformational dynamics that diversify the function of the core-structure to yield transcription factors, enzymes or extra-cytoplasmic transport related proteins. To understand both the mechanistic distinction of these proteins and their evolutionary process, we used single-molecule Förster resonance energy transfer (smFRET)^58^ and Hydrogen Deuterium Exchange Mass Spectrometry (HDX-MS)^59^. smFRET allows to monitor Tier-0 dynamics with a temporal resolution down to microseconds and sub-nanometer spatial resolution at the single-molecule level, even for highly-heterogenous conformational ensembles.^12,60–62^ HDX-MS complements smFRET, as it can probe Tier-1 dynamics throughout the structure that occur within the structural states identified by smFRET (Fig. 1A).^63^ Using this combination of techniques, we show how the C-terminal extensions confer distinct Tier-0 dynamics to the structural core in a manner specific to their three-dimensional orientation. This arrangement also dictates specific geometrical criteria, crucial for establishing specific ligand interactions, all of which we detail in this work. On the other hand, we reveal the means by which N-terminal domain additions enable oligomerization to provide distinct quaternary dynamics in LysR-type transcriptional regulators. The remarkable modularity of these proteins permits us to confirm and expand the evolvability theory for the primordial core-structure during long-period evolution, which is largely facilitated, seemingly, by genetic recombination events.

## Results

### Structure, classification and evolution of the selected bilobed proteins

In this study, our focus was on proteins composed of two globular lobes (bilobed), each of 3 layers (α/β/α), articulated around a central β-sheet hinge (Fig. 1B). This focus meant that not all multidomain architectures harbouring the type-II PBP domains were included. To investigate the evolution of structural dynamics in the selected proteins, we constructed phylogenetic trees based on both sequence and structural information (Fig. 1C). This analysis indicated that the proteins evolved from a common ancestor that diversified into seven distinct structural classes A to G (Fig. 1C/D), members of which are found throughout all kingdoms of life with some even present in viruses (Fig. S1A, SuppFile. 1, 2). They have a consensus structure (Fig. 1B, top row in Fig. 1D and SuppFile. 3), which we have dubbed the *cherry-core* (hereafter *CC*, and proteins harbouring this core, *cherry-core proteins*, *CCPs*) because of the bilobed structure’s resemblance to a cherry (Fig. S2A), that contains the type-II PBP domains.^64^ According to ECOD database, the two domains of *CCPs* belong to the same X-, H- and T-groups (Periplasmic Binding Protein-like II; SuppFile. 3), supporting the common ancestry.^38^ In the consensus structure (Fig. 1C/D), domains D1 and D2 adopt a face-to-face mirror-fashioned geometry with the active site, which is typically a ligand-binding site, located at their interface (Fig. S2B).

The majority of the selected proteins have distinct segments, N-terminally linked to the *CC* (Fig.1C/D, Fig. S1). Class A proteins have the addition of a LysR winged helix-turn-helix (HTH)-type DNA binding domain (ECOD: X-, H-group: HTH, T-group: Winged; Fig. 1C). This element has been shown to be responsible for oligomerization and binding to promoter DNA.^65–67^ Most proteins of classes B-D, and F, G contain N-terminal localization signals for export via the general secretion (Sec) pathway or the twin-arginine translocation (Tat) pathway.^68–71^ The presence of the various address tags was obtained from UniProtKB^72^ and verified manually by inspecting all 600 protein sequences with PRED-TAT.^73^ Class E proteins are predominantly cytosolic and lack such an N-terminal signal peptide.

In addition, classes B-G have distinct C-terminal structural embellishments, hereafter termed C-tails (Fig. 1D). For example, Helical-tail 1 (H1) is common to all classes, except A and C, and importantly, it has a similar placement in the 3D architecture of the proteins. All the other C-tails are unique to a specific class, such as the Helical-tail G (HG), present only in class G proteins.

### Function, ligand specificity and conformational states of the selected bilobed proteins

The function or ligand specificity of the *CCPs*, as documented in UniProtKB^72^, was found to correlate with the assigned structural class (Fig. S1A). Class A proteins are bacterial transcription factors of the LysR-type transcriptional regulator (LTTRs) family. Class E proteins are predominantly eukaryotic single-turnover enzymes. The majority of the remainder (class B-D, F, G) are found in prokaryotes, and associate with the translocator domains of ABC transporters, or with the membrane-embedded domains of chemoreceptors, where they mediate unidirectional solute transport and signal transduction respectively.

To investigate what role the distinct C-tails of these proteins might have played in the evolution of structural dynamics and function, we examined *CCPs* for which high-resolution structures of unliganded (*apo*) and liganded (*holo*) states were available (Fig. 2, Table S1). Interestingly, class A and E proteins, in the majority of cases, display nearly identical *apo* and *holo* structures. They also show the widest variety of substrates with little chemical structure similarity (Fig. S3). For most members of the other classes, D1 and D2 of the *CC* undergo a rigid body rotation of varying degrees (Fig. 2, Table S1). For the solute binding proteins this mode of substrate binding has been called the “Venus-Fly Trap mechanism”.^74^ These proteins also recognize ligands with a specific pharmacophore^75^: amino acids, ethanolamines, phosphonates, iron-phosphate complexes and carbohydrates are recognized by classes B, C, D, F and G, respectively (Fig. S3). Another striking difference is that proteins in classes B-G are monomeric, whereas those in class A are oligomeric (see sections below).

**Figure 2.**
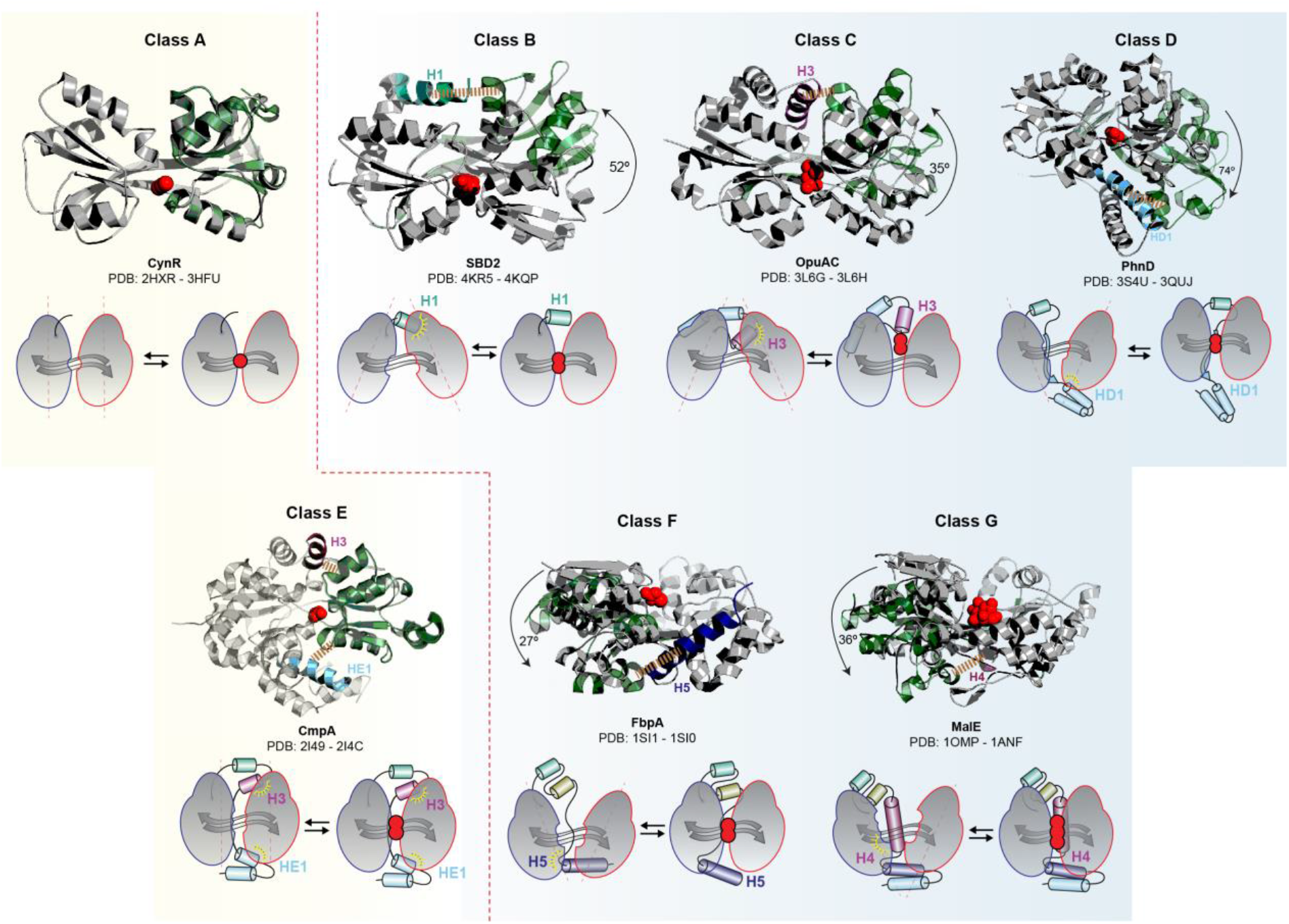
Structures of *CCPs* highlighting their C-tails. Structures (top) and schematics (bottom) of the identified structural classes (A-G). The *apo* and *holo* structures of the indicated *CCPs* are superimposed to highlight the role of the C-tail. The domain to which the two structures were superimposed is represented in grey (D1 in A-E and D2 in F, G), whereas the one that is displaced relative to the C-tail in the open state is represented in green. Arrows and schematics indicate domain displacement (see also Table S1). Interaction contact maps are presented in Table S2. The most prominent contacts between the indicated secondary structure elements of the C-tail and D1 (F, G) or D2 (B-E) that stabilize the open state are indicated by a yellow dashed line. The PDB codes and protein names are indicated.

### smFRET to monitor large domain motions (Tier-0) in bilobed proteins

To investigate structural changes and dynamics of the *CCPs*, we used smFRET^12,60–62^ to probe Tier-0 dynamics at near-physiological conditions in aqueous buffer at room temperature. The following representative proteins (details in Fig. S1) were investigated: the effector binding domain (EBD) of CynR, representing the *CC* of full-length CynR^66^ (class A), SBD2^76^ (class B), OpuAC^77^ (class C), CmpA^78^ (class E) and MalE^79^ (class G). For these experiments, D1 and D2 were stochastically labelled using cysteines that were substituted for non-conserved, surface-exposed residues, one in each domain (Fig. 3A). Fluorophore labelling was performed by maleimide-thiol conjugation.^22,80^ Labelling positions were selected based on the crystal structures, and the residues chosen display large changes in separation between the *apo* and *holo* states. smFRET was performed by confocal microscopy with alternating laser excitation (ALEX).^60^

**Figure 3.**
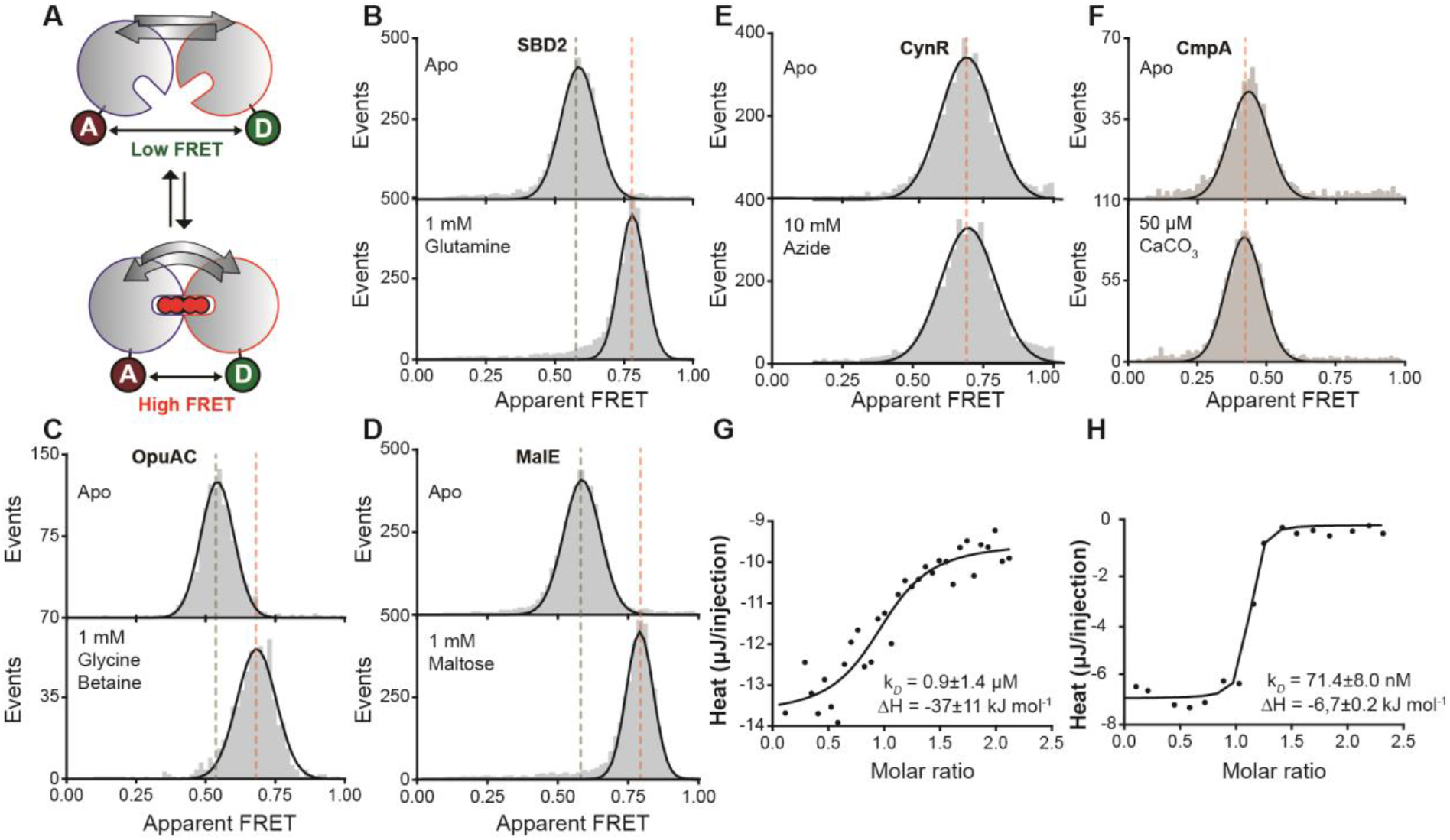
Monitoring structural changes and ligand binding in *CCPs* using smFRET and ITC. (**A**) Schematic of experimental strategy to monitor structural states by FRET efficiency via stochastic labelling of D1 and D2 with donor and acceptor fluorophores. (**B**-**F**), Solution-based apparent FRET efficiency histograms in the absence (top) or presence (bottom) of saturating ligand concentrations for each protein indicated. Grey bars are experimental data and solid line is the fit. Centre position of Gaussian fits are given in Table. S5. (**G**, **H**) Binding isotherms of the calorimetric titration of azide (G) and calcium-carbonate (H) to CynR (G) or CmpA (H), respectively with the indicated thermodynamic parameters. According to high-resolution structural data^84^, both Ca^2+^ and CO_3_^−^ are present in the CmpA binding cleft. Indeed, we observed heat release upon titration of Ca^2+^ to a CO_3_^−^ bound CmpA. Data points represent the heat of reaction per injection and the line is the fit.

As predicted by our structural analysis (Fig. 2, Table. S1), and in line with our previous observations^81^, the FRET efficiency histograms and fitted distributions (Fig. 3B-D) shifted towards higher FRET efficiency (*E*) values upon addition of saturating concentrations of ligand, for SBD2, OpuAC and MalE. This indicates that in the *apo* state, the donor and acceptor dyes are further apart (Fig. 3B-D upper panels, low FRET) compared to the *holo* state (Fig. 3B-D, lower panels, high FRET). Thus, our data suggest that ligand binding drives Tier-0 dynamics in these *CCPs*.

In contrast, the distributions of class A and E proteins, CynR^82^ and CmpA^78^ respectively (Fig. 3E, F), were virtually identical in the absence or presence of saturating ligand concentrations. Ligand binding was confirmed via isothermal titration calorimetry (ITC) showing binding affinities of both proteins in the micromolar range (Fig. 3G, H). Thus, we conclude that the *CC* of class A and E proteins lack Tier-0 dynamics on the probed reaction coordinates for the selected FRET-distance pairs in contrast to the other structural classes.

Interestingly, this observation is in line with the known biological function. Class E proteins are predominantly single turnover enzymes (Fig. S1A), for which the rigidity of their active site is a prerequisite for catalysis^83^ (Fig. S4A). As previously suggested by us and others, Tier-0 dynamics in periplasmic-binding proteins (class B, C, D, F and G) are utilized in the regulation of transport in ABC importers (see discussion). A remaining question is, however, how ligand binding in class A proteins triggers transcriptional processes without major structural changes related to ligand binding.

### HDX-MS to study local flexibility (Tier-1) of a class A protein, CynR

The DNA binding domain of class A proteins typically comprises a ~58 aa HTH motif. This is followed by a ~20 aa long helix that provides a dimerization interface and a connecting loop that attaches the DNA binding domain to the *CC* (Fig. 4A). The *CC* acts as the tetramerization interface within the full-length *CCP* or the dimerization interface within the effector binding domain (Fig. 4B, Fig. S4B-E). Indeed, by using SEC-MALS we observed that (full length) CynR is tetrameric, whereas the *CC* of CynR is dimeric (Fig. 4C).

**Figure 4.**
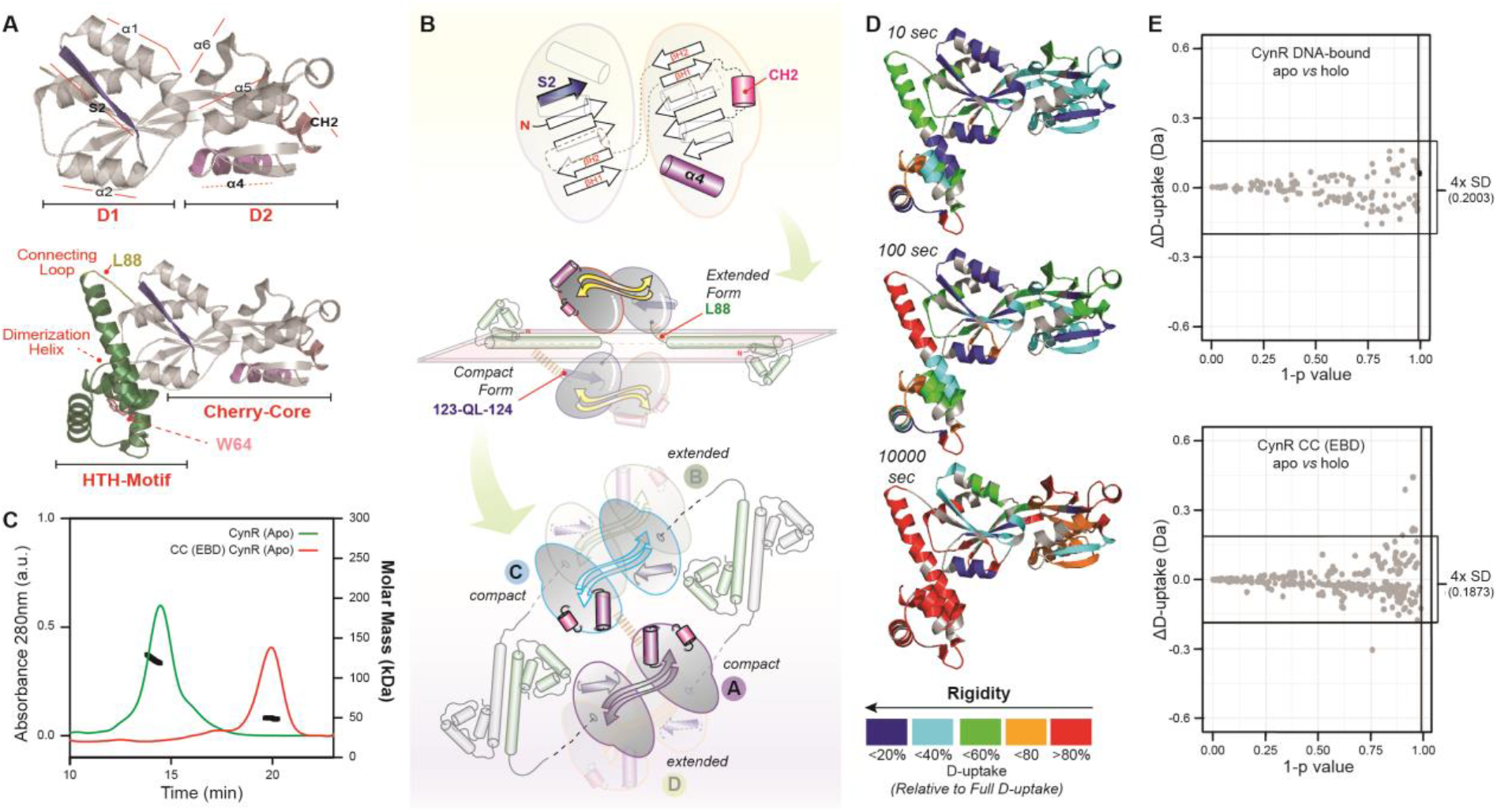
CynR tertiary and quaternary assemblies probed by MALS and HDX-MS. (**A**) Crystal structure of the CynR *CC* (PDB: 2HXR) with colored secondary structural elements discussed in the text that are critical for its quaternary dynamics (top panel). A homology model of full-length CynR obtained from the SWISS-MODEL server with CbnR (PDB: 1IZ1) as a template (bottom panel). The HTH domain is colored green and the loop connecting it to the *CC* splitpea. W64 at the tip of the dimerization helix and L88 of the connecting-loop are indicated with an arrow. (**B**) top panel: Schematic representation of the *CC* of CynR using the same color-coding as in A. middle panel: Schematic representation of the CynR homology model, with one of the protomers in the compact and the other in the extended configuration. The *CC* of CynR protomers self-associate to form dimers, or for the full-length protein, tetramers. For clarity, two of the protomers have been omitted. Bottom panel: The tetrameric assembly formed by the self-association of the dimerization helices and the *CC*. Of the two interacting protomers, one is present in the compact configuration, whereas the other is extended. The two protomers with the compact configuration are shown at the top of the plane, whereas those that are extended are at the bottom. (**C**) SEC-MALS analysis of full-length and *CC* of CynR (3 μM). UV traces of the chromatograms were superimposed on the measured mass (black cycles). (**D**) Structural dynamics of full-length CynR in the absence of ligand and DNA by HDX-MS. Deuterium uptake values are reported for the incubation times in deuterated buffer and expressed relative to the fully deuterated control. These values are mapped onto the CynR homology model (as in A), using the indicated color gradient. Proline residues as well as the first residue of each peptide were excluded from mapping, as they do not contribute to the observed D-uptake. (**E**) Scatter plot visualization of the statistical analysis of D-uptake differences between the apo and holo states of full-length CynR (upper panel) and the *CC* of CynR (bottom panel). Three statistical criteria were used (Materials and Methods), as described previously.^89^ Statistically significant differences would appear as black spheres (indicating that ΔD-uptake>2xSD for a specific peptide), lying outside the 99% confidence threshold (1-p≥0.99; indicated on y-axis) and outside the ± 4 x pooled average SD cut-off (indicated on x-axis; value given on the right).

To determine whether ligand binding to CynR resulted in conformational changes that were not detectable along our selected reaction coordinate, or that were too small or fast for smFRET (Fig. 3E), we performed HDX-MS. In contrast to smFRET, which reports on a single distance along a single reaction coordinate, HDX-MS probes structural dynamics at near-residue level resolution providing global insights into Tier-1 dynamics. HDX-MS detects the exchange of hydrogens with deuterium at solvent-accessible and non-hydrogen-bonded backbone amides.^85,86^ Hydrogens involved in stabilizing the secondary, tertiary or quaternary structure of a protein via hydrogen bonds are exchanged more slowly through structural transitions that disrupt these bonds. Deuterium incorporation into the protein can then be determined, following proteolysis, by mass spectrometry. The mass difference between hydrogen (^1^H) and deuterium (^2^H) results in a mass shift between non-deuterated and deuterated peptides that is a measure of the number of exchanged hydrogens.^59,87,88^

For such investigations, the *CC* of CynR or full-length CynR were isotopically labelled (pD, 7.4; 25 °C) for different time periods (10 – 10^5^ sec) either in its free or in its DNA bound state, and the peptic fragments were identified by MS (Fig. 4D, SuppFile. 4). For each peptide, the fraction of deuterium uptake relative to the maximum determined deuterium incorporation was calculated (SuppFile. 4). The data reveal a rigid character of the *CC* of CynR, whereas the dimerization helix and the rest of the DNA binding domain turned out to be more flexible (Fig. 4D). To identify the regions in which pronounced structural changes were induced by azide binding, we performed comparative HDX-MS. For this, we determined the difference of deuterium uptake (ΔD) for each peptide between different conditions, e.g., CynR *apo* versus *holo* states. Observed differences would indicate a decrease (protection) or an increase (deprotection) of deuterium uptake upon azide binding (SuppFile. 4). No statistically significant change in the deuterium uptake was observed upon azide binding (Fig. 4E). From this, we can conclude that no detectable change in structural dynamics is induced by the ligand in the *CC* of CynR in its free or DNA bound form (Fig. 4E, SuppFile. 4). This includes Tier-0 dynamics (detected by smFRET and HDX-MS), Tier-1 or quaternary dynamics (both detected by HDX-MS).

### An asymmetric C-tail drives large-scale tertiary (Tier-0) structural changes

From our results on the selected *CCPs*, only those with a C-tail, display Tier-0 dynamics (Fig. 2, 3). To investigate how the C-tails introduced Tier-0 dynamics to the *CC*, we compared crystal structures of the *apo* and *holo* states of the *CCPs* to identify interactions between their C-tails and the *CC*. Fig. 2 summarizes the results of these comparisons. Interestingly, we found that the *holo* structures are similar for all *CCPs*, but the *apo* structures are class-specific. Contact mapping of the interactions between the *CC* and C-tail using the protein interaction calculator web-server^90^, showed that the number and characteristics of the interactions depends on the structural class (Fig. 2, Table. S2). Strikingly, these interactions predominantly stabilize the open conformation of the *CC*, the only exception being for the class A and E. In the former cases, stabilization is associated with an asymmetrical placement of the C-tail with respect to D1 and D2. In contrast, the C-tail of class E (predominantly H3 and HE1; Fig. 2) is placed symmetrically around D1 & D2, and thus cannot provide the required structural asymmetry needed to create a stable open state. The stabilizing interactions of the C-tail to form the open state involve the consensus *CC*-helices of D1 and D2 (Fig. 2, Table. S2). In classes B, C and D such interactions involve D2, whereas D1 is contacted in classes F, and G. These asymmetrical interactions create active sites with distinct geometries (Fig. 2, Fig. S2).

In alternative phylogenetic trees, based on the protein sequence using either D1/D2-domains or the C-tail, the clustering remains similar (Fig. S1B-D, SuppFile. 1, 2); indicating that D1/D2 (*CC* domains) and the C-tail of a specific class co-evolved to be part of the same polypeptide. This is in line with the role of the C-tail to interact with one specific domain, D1 or D2, to stabilize the open structural state.

### Experimental verification of the role of C-tail interactions in stabilizing the open conformation

To confirm the role of the C-tail interactions with the rigid domains D1 or D2 for stabilization of the open state, we manipulated the relevant ones (Fig. 2; Table. S2). The impact was tested via assessing structural states and ligand binding affinities, monitored in smFRET experiments. Test cases were selected from class B and G, as these *CCP* classes are only remotely related by the first major clade in the evolutionary trees (Fig. 1C) and their open state is stabilized by distinct helices within the C-tails (Fig. 2), contacting either D2 (class B) or D1 (class G).

In SBD2, a hydrophobic interaction between L480 in the C-tail and P419 in D2 was weakened by substitution of L480 with alanine (L480A; Fig. 5A, Table. S2). The mutation resulted in the appearance of a subpopulation of molecules (~15%) that were in the closed state in the absence of glutamine (Fig. 5C *vs* B). To rule out the possibility that this might have been due to an artefact introduced by the choice of fluorophores, a second pair of fluorophores was tested, which showed a comparable result (Fig. S5). We also examined whether residual endogenous ligand might account for this subpopulation by performing smFRET measurements on diluted samples, but these experiments showed subpopulations of a similar size (Fig. S5). A closed-unliganded conformation was also observed for the wild-type SBD2 previously, but with a much lower abundance (~1%). Detection of this small subpopulation required the use of scanning confocal microscopy^81^, because populations <5% cannot be detected reliably with ALEX microscopy. We also observed small differences in the mean *E* values for *apo* and *holo* states of SBD2 (L480A) as compared to SBD2 suggesting that the structural landscape had been altered (Fig. S5). Destabilizing the open state is supposed to decrease the K_d_ ^91^, consistent with a 2-fold increase in its glutamine binding affinity of SBD2 (L480A) as compared to the wild-type (K_d_ of 840 ± 270 nM and 1990 ± 130 nM, respectively; Fig. 5D).

**Figure 5.**
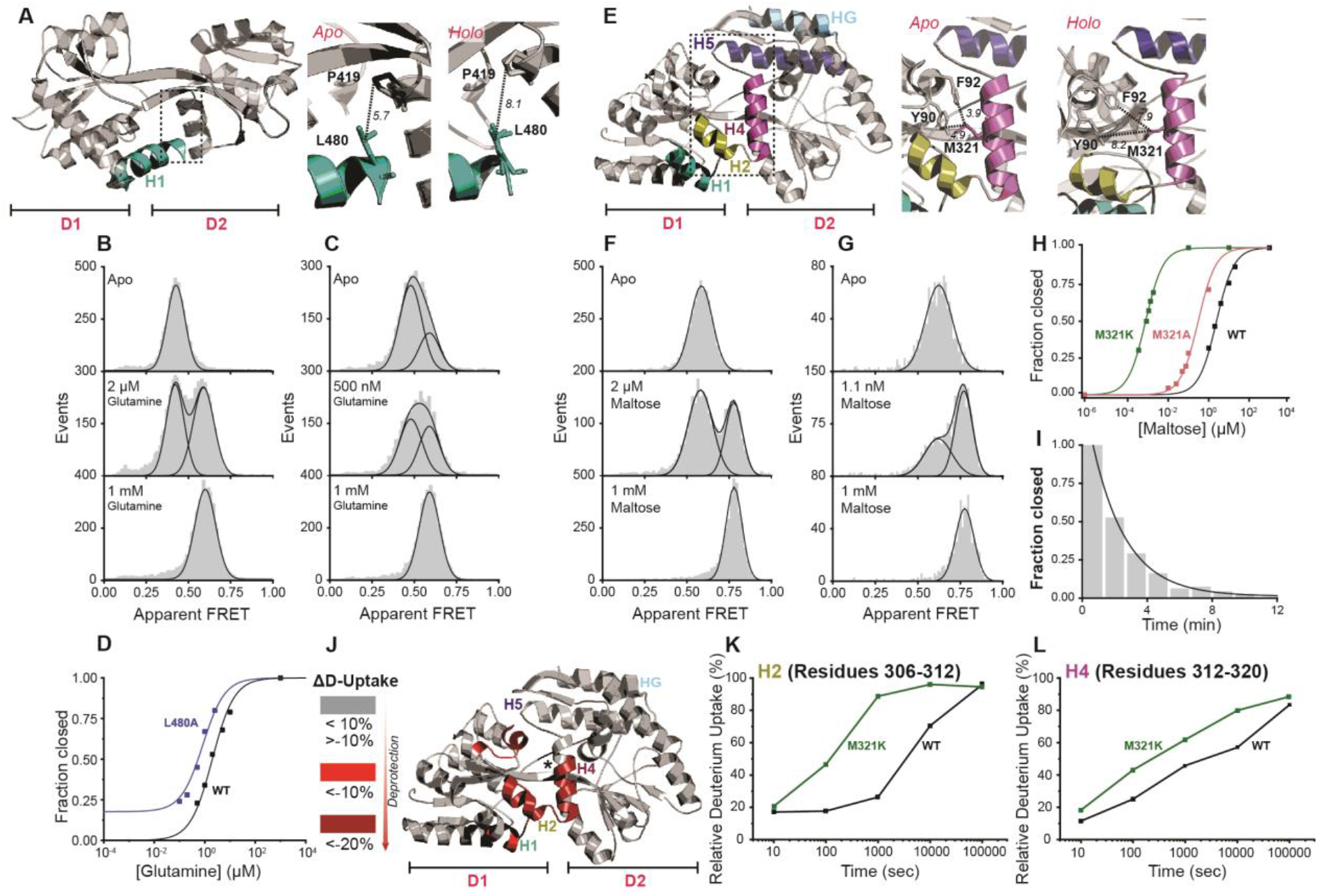
Experimental verification of the role of C-tail interactions in stabilizing the open conformation of SBD2 and MalE by smFRET and HDX-MS. (**A**, **E**) Dotted rectangles (left panels) on the SBD2 (in A PDB: 4KR5) and MalE (in E PDB: 1OMP) structures highlights the critical contact region between the C-tail and the *CC* that stabilize the open state. Zoom in of rectangle regions (middle and right panels) depicting interactions between the C-tail helix H1 and D2 (A) or between the C-tail helix H4 and D1 (E) in the indicated *apo* (open) or *holo* (closed) states. Distances (Å) between L480 and P419 (A) or between M321 and Y90/F92 (E) are shown as black dotted lines. (**B**, **C**) Solution-based apparent FRET efficiency histograms of SBD2 (B) and SBD2 (L480A) (C) at different conditions as indicated. (**D**) Fraction of the closed state (high FRET state; Fig. S5) of SBD2 and the indicated derivative as a function of glutamine concentration. (**F**, **G**) Solution-based apparent FRET efficiency histograms of MalE (F) and MalE (M321K) (G) at different conditions as indicated. (**H**) Fraction of the closed state (high FRET state; Fig. S6) of MalE and its derivatives as a function of maltose concentration. (**I**) Maltose release from MalE (M321K) over time determined by solution-based smFRET (see detailed values in Figure S6). Data points (panel D, H) and grey bars (panel B, C, F, G and I) are the experimental data, and the solid line is the fit. **(J)** Map of regions in MalE structure that show statistically significant increase in deuterium uptake caused by M321K (numerical values and complete statistical analysis presented in SuppFile. 5). n=3. (**K, L**) Deuterium uptake for the indicated MalE C-tail helices. Standard Deviations (SD) are shown in the deuterium uptake plots.

For MalE, we constructed the derivatives MalE (M321A)^92^ and MalE (M321K), with weakened interactions between the C-tail H4 and D1, by disrupting the hydrophobic interaction of M321 with Y90 and F92, and the aromatic sulphur interactions with Y90 (Fig. 5E, Table. S2). For MalE (M321A), the stabilizing interactions of M321 are partially abolished. This resulted in an 8-fold increase of maltose affinity [*K*_*d*_ from 2400 ± 400 nM in MalE to 300 ± 50 nM in MalE (M321A) (Fig. 5H, Fig. S6)]. An even stronger effect was observed for MalE (M321K) with an affinity enhanced by 3000-fold (*K*_*d*_ of 0.81 ± 0.15 nM; Fig. 5G *vs* Fig. 5F, Fig. 5H). Shilton and co-workers have also proposed the hydrophobic interactions of M321 as being important structural determinants of the open state^92^, affecting the affinity of MalE for maltose, in agreement with our results.

We next compared the lifetime of the closed maltose-bound conformation of MalE (M321K) with that of wild-type. Addition of 10 nM maltose allowed MalE (M321K) to occupy the closed state exclusively (Fig. S6I, top panel). 20 µM unlabelled MalE (M321K) protein was subsequently added to scavenge maltose, which is stochastically released from the labelled protein. In a time-course experiment the decrease in the population of closed maltose-bound MalE (M321K) was then followed as a function of time^93^ (Fig. 5I, Fig. S6I). From these experiments, we found that the lifetime of the closed maltose-bound conformation of MalE (M321K) was 122 ± 12 s, and that this value was 2500-fold higher that the value determined for wild-type MalE (0.048 ± 0.010 s; Fig. S6). This result is consistent with the observed increase affinity of the derivatives for maltose. Our observations clearly demonstrate that the C-tail is crucial for the conformational landscape of the *CC* and the modulation of the states by ligands.

### The C-tail dynamics control the structural transitions of the *cherry-core*

In the above-mentioned derivatives of SBD2 and MalE, the structural basis for the destabilization of the open state could not be addressed mechanistically. Our smFRET experiments report predominantly on Tier-0 dynamics that affect the protein’s tertiary structure (Fig. 1A). Considering that Tier-0 dynamics, as a consequence of C-tail-D1/D2 destabilization, could only be observed for SBD2; we performed comparative HDX-MS (SuppFile. 5) to monitor changes in the secondary structure of MalE in comparison to MalE (M321K), (Fig. 5J). For this, we determined the difference of deuterium-uptake (ΔD) for each peptide between MalE and MalE (M321K) in their *apo* states. Remarkably, comparative HDX-MS indicates that the differences between wild-type and MalE (M321K) are localized almost exclusively at the C-tail and specifically in regions interacting with D1 (SuppFile. 5, Fig. 5J). The D-uptake of the C-tail helices H2/H4, that are critical for stabilizing the open state (Fig. 5E, Table. S2) denotes that their rigidity was significantly reduced in the MalE derivative (Fig. 5 K, L). As might be expected, reduced rigidity occurred also in a region within the *CC* containing Y90 and F92. Notably, the same regions become allosterically destabilized in wild-type MalE upon maltose binding (SuppFile. 5). These results support the idea that the mutation leads to a weaker interaction between the C-tail and the *CC* resulting in a destabilization of the open conformation, because these C-tail elements and the region containing contact residues were found to be more flexible and solvent-exposed.

Taken together, these results provide compelling evidence that the interactions between the *CC* and C-tails are involved in stabilizing the open conformation of the *CC* and thus evolved to stabilize a novel conformational state.

## Discussion

Common (structural) origin represents the hallmark of Darwinian evolution. Homology, or descent from a common ancestor is often deduced from similarities in protein sequences or better from structures, as the latter are more conserved during evolution.^94^ However, similar structures can originate from divergent, convergent or parallel evolution.^95^ The most common “tricks” nature uses to vary a protein domain are: β-strand invasion/withdrawal, insertions/deletions/substitutions of secondary structure elements, domain flip/swaps and circular permutations.^96,97^ The currently established evolvability theory relies on investigations involving a fixed length polypeptide chain by observing the effects of sequence variations, accomplished primarily by directed evolutionary approaches or by investigating closely related functional homologues. The functional promiscuity originating from structural variability is altered by the few amino acid modifications that can yield alterations of local structural fluctuations. For this reason, this theory can well explain protein evolution during short time periods.

In this study, we focused on the analysis of structures that have diverged over longer evolutionary periods. We analyzed a group of ∼600 proteins that share a core structure with the same topology of secondary structure elements giving rise to identical three-dimensional structures. The structure was predominantly varied by terminal embellishments and exhibits detectable sequence identity, used for constructing the sequence-based phylogenetic trees (Fig. S1). The identified proteins show divergent long-term evolution from a common ancestor, which spreads throughout the tree of life. This common ancestor is represented by the consensus core structure (dubbed *cherry-core*, *CC*, Fig. S2), and encountered within the type-II class of periplasmic binding proteins.^64,98^ As proposed previously^64^, the *CC* derived possibly from a gene duplication of a type-II periplasmic binding protein domain [(ECOD); SuppFile.3] that has been connected by two β-strands (Fig. 6A).

**Figure 6.**
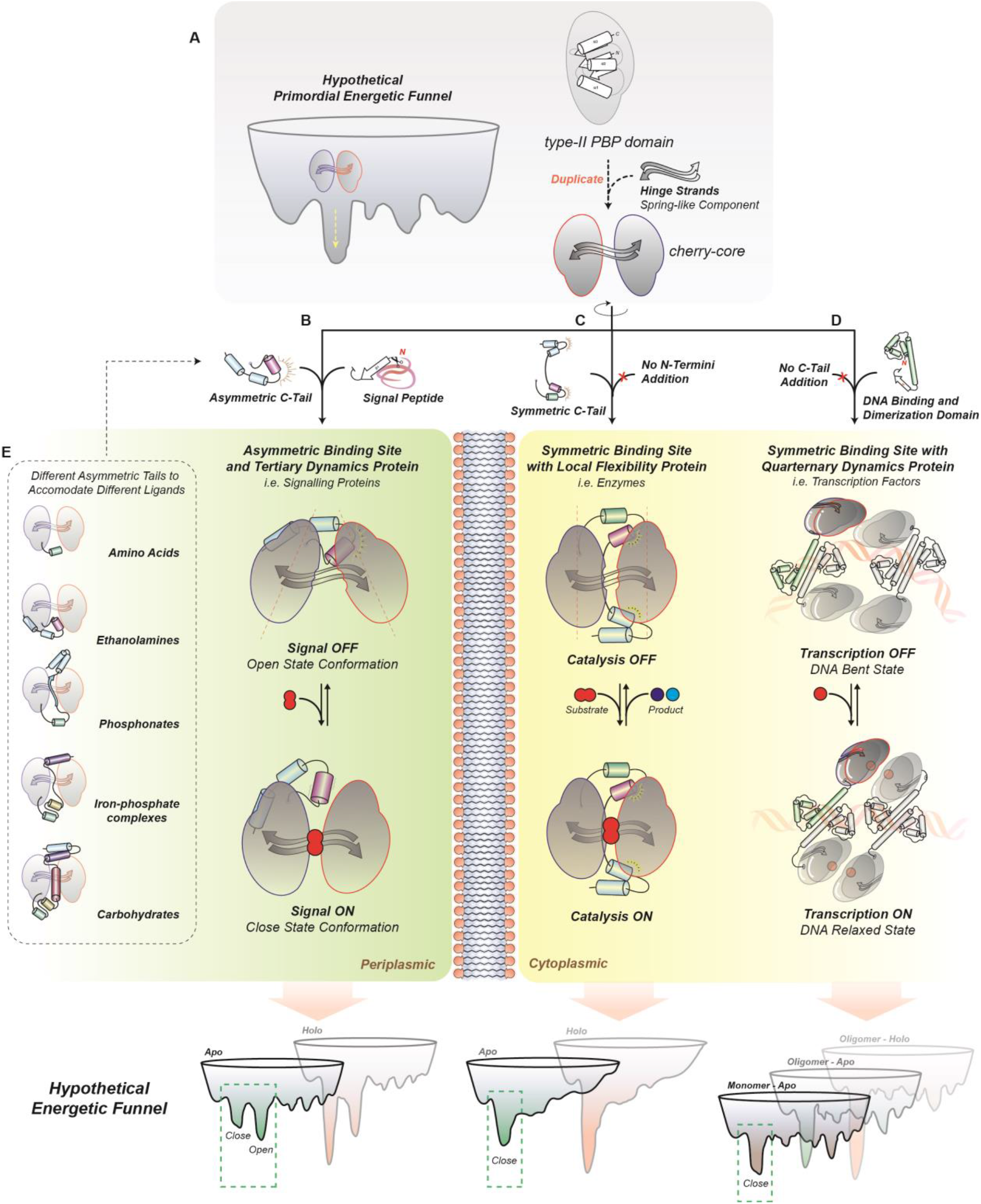
Model for the evolution of the *cherry-core* and hypothetical energetic funnels: A gene duplication of a type-II periplasmic binding protein domain connected by a single β-sheet gave rise to the *CC*, being composed of a unique closed structural state, represented by a single-well in the energetic funnel. The function of the *CC* was uniquely binding. Extant proteins acquired different extensions that altered their localization and dynamics and by that their function and specificity. An N-terminal signal peptide and an assymetrical C-terminal tail generated extra-cytoplasmic proteins evolving an additional open state. The different flavors of open states, conferred distinct substrate specificities. The 2 states of *CCPs*, predominantly, signal substrate transport by their association to the membrane-embedded translocator domains of (ABC) transporters. A symmetrical C-tail rigidified the closed state and yielded primarily single-turnover enzymes. The N-terminal domain addition of a flexible dimerization helix and a DNA binding motif (HTH-type) conferred distinct oligomeric assemblies with different quaternary dynamics, yielding transcription factors. For details see text and Fig. S8.

### *CCPs* with asymmetric C-tails and N-terminal signal peptides

When the *CC* is fused N-terminally to a signal peptide and C-terminally to an asymmetric C-tail, the *CC* acts as an extra-cytoplasmic monomeric protein, that associates with the translocator domains of ABC transporters or with chemoreceptors (Fig. S1, Fig. 6B). The two structural states are *apo*-open *vs. holo*-closed and originate from Tier-0 dynamics, which are critical determinants of their biological function: the membrane-embedded partners can discriminate between open *vs* closed states to activate or inactivate a biological process^80,99–102^, such as solute transport. These structural transitions rely on the “generation” of an open state of the *CC*, accomplished by the C-tail, through its asymmetric interactions with either of the domains of the *CC* (D1 or D2, Fig. 2). Evidently, the open state is stabilized by such enthalpic contributions and destabilized by mutations or ligand binding that increase the flexibility of the interacting regions (Fig. 5J) [the C-tail elements with either D2 (classes B, C, D) or D1 (classes F, G), (Fig. 2)]. Seemingly, entropic contributions (protein conformational entropy) bias the structural equilibrium towards the closed state. Given the fact that the *holo* is the closed state driven by ligand binding^81^, we anticipate that the interactions of the ligand with the *CC* cleft allosterically induce order-to disorder transitions to alter the structural equilibrium. Depending on the placement of the asymmetric C-tail in the three-dimensional space, we identified five different “flavors” of open states, establishing distinct geometries of active sites; all resembling a triangle distinctly oriented in space (Fig. 2, Fig. 6E). We verified that each active site geometry can recognize a specific chemical structure (Fig. S3), in full agreement with ligand binding triggering closing.

The evolvability theory of Tawfik and co-workers^29,51^ explains the functional promiscuity encountered within the different structural classes. We selected two closely related proteins in our phylogenetic tree in class B (Fig. 1C) to illustrate this (SuppFile. 2, red asterisk). SBD1 (PDB: 4LA9) mediates the unidirectional transport of two substrates (glutamine & asparagine), whereas SBD2 (PDB: 4KR5) transports one of them (glutamine), though captures it with higher affinity.^80^ SBD1 can be transformed to bind glutamine with higher affinity alike SBD2 by mutating three amino acids.^76^ Clearly, functional promiscuity is traded seemingly with ligand specificity.^56^ However, only the modularity notion introduced in this study can explain a greater ligand-functional promiscuity. Class B proteins recognizing and mediating amino acid transport (Fig. S1), would gain tertiary structural variability by acquiring four more helical elements at their C-termini (H2, H4, H5, HG, Fig. 1C, D) (Fig. 6). This would potentially allow to switch them to class G, to bind and mediate transport of carbohydrates (Fig. S1A). According to another evolutionary trajectory, class B proteins, by acquiring helices H3, HE1, HE2; could divert to class E, and switch from transport related proteins to enzymes (Fig. 6) by restricting their tertiary structural variability to a unique closed state. An extreme case of functional divergence has been experimentally verified in a recent study that restored the evolutionary history of the enzyme cyclohexadienyl dehydratase^103^, a class B member according to our classification. The reconstructed ancestor of this enzyme was a highly promiscuous transport-related protein possessing the open-unliganded and the closed-liganded structural state and was able to bind four different cationic amino acids. On the other side, cyclohexadienyl dehydratase binds a single substrate and forms a unique closed state for its catalytic function, since it is believed that structural sampling represent a constrain for catalytic activities.^104,105^ Our study can now explain the means by which the addition of modular elements to a conserved core-structure diversify its function, i.e., by modulating its structural landscape.

### *CCPs* with symmetric C-tails

When the *CC* is combined C-terminally to a symmetric C-tail during evolution (Fig. 6C), the *CC* either operates as an enzyme or is associated with ABC transporters (Fig. S1). Those proteins are present in a unique (*apo* and *holo* state) closed structure (Fig. 2), as asymmetrical interactions are an essential prerequisite for open state formation. Interestingly, these ABC transporter-associated proteins are both interacting with the actual translocator from the extra-cytoplasmic side (alike the classes B, C, D, F, G; described in the previous paragraph and in such a case possess also an N-terminal signal peptide), but are also tethered covalently to the translocator ATPase motor domains.^84^ As the switching (open to closed) behaviour – to activate the transport cycle – is missing^80,81,99^, the non-canonical arrangement of these ABC transporters could only trigger transport activation using a yet uncharacterized mechanism. On the other hand, the lack of Tier-0 dynamics observed in the single-turnover enzymes is conforming to their enzymatic mechanism demanding an extremely rigid active site (Fig. S4).^83^ Evidently, the rigidity required for the chemistry is so dramatic that the flexibility of a non-covalently (to the polypeptide chain) bound histidine to the cleft would render the reaction unproductive.

### *CCPs* with N-terminal domains

Lastly, the *CC* bearing an N-terminal domain harboring the HTH type DNA binding domain, but no C-tail (Fig. 6D), yield transcription factors of the LTTR family. Apparently, in such a case, Tier-0 dynamics of the *CC* are not required for function (Fig. 3E). The rigidity of the *CC* (Fig. 4D) is evidently required for the stability of the quaternary assemblies, as those have large cavities and holes (Fig. S7).^106–111^

### Structural basis of the transcriptional activation by the LysR-type regulators

Our data and analysis gave insights into the steps taken during evolution to shape the *CC* into functional ligand-receptor associated proteins (classes B, C, D, F, G) but also one-turn enzymes (class E). Yet, how the ligand-driven signal propagates within the LTTR family, such as CynR, remains largely elusive.

The *CC* of CynR constitutes only one part of its structure, i.e., the sensory effector-binding domain (EBD), of a full-length transcription factor of the LTTR family with an additional winged helix-turn-helix (HTH)-type DNA binding domain (Fig. 4A/B). Although only little direct experimental evidence is available, it has been proposed that changes in the tetrameric assembly of LysR-type transcription factors can be induced by ligand binding.^66,112^ These quaternary dynamics were suggested to activate transcription by a transition from a bent DNA (transcription OFF) to an unbent state (transcription ON)^66,112^. Our smFRET and HDX-results (Fig. 3E, Fig. 4E), however, did not reveal any detectable changes of the *CC* upon azide binding. This suggests that ligand-driven structural changes within the *CC* protomers are minor and might thus be very hard to detect.

To gain insights into the quaternary dynamics that govern function, we analysed the available high-resolution structures of the oligomeric assemblies belonging to the LysR transcription factors^106–111^; to identify the evolutionary relevant ones by EPIC.^113,114^ Subsequently, we modelled the CynR sequence after these structures in the presence or absence of DNA (Fig. S7). The structural basis of the distinct tetrameric assemblies is the positioning of the dimerization helix with respect to the *CC*, dictated by the connecting loop (Fig. 4A/B). In the extreme case that an additional helix (RD1-CH5, see Supp. File 3) is linked to- and displaces the connecting loop, an octameric assembly is obtained (Fig. S7H). To shed light on this, we monitored intrinsic Trp fluorescence during thermal melting (Table. S3). From the two tryptophanes of CynR; one present in the dimerization helix (aa 64) and the other on D1 (aa 274) of the *CC* (Fig. 4A, B); only the former one contributes to a fluorescent signal (data not shown). Under DNA-free *apo* conditions, CynR displays a T_*m(app)*_ ~ 55°C that is significantly destabilized (~6°C) after binding to DNA. Addition of azide at free or DNA bound CynR causes a ligand-characteristic signature throughout the Trp-temperature spectrums, giving rise to a secondary T_*m(app)*_ ~ at 27°C in both cases (Table. S3).

To understand the signal propagation originating from ligand binding, we inspected the available structural information (Fig. S7). In agreement with the experimental findings (Fig. 3E, Fig. 4E), our structural analysis indicated that D1-D2 motions are not required (Table. S4) for the oligomeric assemblies of the same transcription factor (OxyR) to differ dramatically in order to trigger structural changes on DNA (Fig. S7E-G). What varies between those assemblies is the order-disorder of specific secondary structure elements within D2 of the *CC* (S4 till α5, see SuppFile. 3). We anticipate that such changes in the flexibility of secondary structure elements are induced by ligand binding, alike in the case of MalE (Fig. 5J-L).

We conclude that signal propagation in *CCPs* comprising large D1/D2 rearrangements (Fig. 2; classes B, C, D, F, G) driven by the C-tail/D1-D2 interactions involves bending/unbending of the “spring-hinge”.^98^ On the other side, propagation in class A is initiated by small/localized rearrangements of secondary structure element within the *CC* somehow transmitted to the N-terminal HTH domain leading to global quaternary structural changes (Fig. S7, S8). Additional analysis regarding the molecular mechanisms of LTTR-transcriptional regulators will be the subject of future studies.

### The energy landscape of the *CCPs* over time

Figure 6 summarizes our proposed evolutionary path taken by the *CC-proteins* over long time periods. The terminal modules that regulate the multi-Tier structural dynamics can be described by the folding funnel model.^8^ The *CCPs* with asymmetric C-tails displaying Tier-0 dynamics (i.e., open-closed states) have the two characteristic wells in the funnel separated by a large energetic barrier (Fig. 6B & Fig. S8C). For that reason, such proteins are found predominantly in the lowest energetic level open state (*apo* energetic funnel; Fig. S8C), with infrequent transitions to the closed one (Fig. 5B, C). Only in the SBD1/2 protein (class B), ~1% occurrence of a closed state has been experimentally observed.^81^ By destabilizing the open state of SBD2 further and shifting it to higher energetic levels, we obtained a ~15% occurrence of a closed state (Fig. 5C). As in the *holo* state energetic funnel, the lowest energy state is the closed one (Fig. S8C); addition of the ligand shifts the equilibrium towards the closed state. Since the dissociation constant (*K*_*D*_) derives from the difference between the lowest energetic levels of the *apo* and *holo* funnels (Fig. S8), destabilization of the C-tail (Fig. 5D, H) that directs the open state of the *apo* funnel at a higher energetic level, leads to an increased affinity.

Alternatively, *CC*-proteins with a symmetric C-tail (Fig. 6C) have a single main energetic well, defining the unique closed structural state both in the *apo* or *holo* funnels (Fig. S8D).

The *CCPs* with N-terminal domain additions (Fig. 6D) have multiple energetic funnels corresponding to oligomerization states (Fig. S8E). The energetic funnel of such *CCPs* is having multiple minima corresponding to the many different arrangements of the flexible N-terminal domain with respect to the *CC* (Fig. S8D). Self-association (oligomer formation) and/or binding to their partners/ligands deepens specific wells that are required for function.

We anticipate the energetic funnel of the primordial consensus *CC* to be extremely rugged, with small energetic barriers between the wells, with a single well somewhat deeper (corresponding to the closed conformation), thus allowing sampling of multiple structural states (Fig. S8A). Such a structural variability would lead to increased substrate promiscuity, however exploiting extremely weak interactions (i.e. low binding affinities; Fig. S8B). The function of the primordial *CC* having no modules could be solely that of binding. In line with the evolvability theory, a specific energetic well of the funnel can be shaped and deepened with multiple rounds of neutral mutations/plastic modifications, trading promiscuity with specificity. The evolutionary pressure for this, is to select the already deeper closed-state well, as its ruggedness (frustration)^115^ can account for high-affinity recognition of very diverse substrates. Addition of a module (internal to the core, or at its termini) can also simultaneously direct the evolutionary pressure to another well specifically, and by that create new functionalities (Fig. 6, Fig. S8): (i) In the case of proteins modulating transport or chemotaxis, an open state well is needed (being much deeper than the closed state well), and different open state wells can be created. The module also has to specifically alter not only the *apo* but also the *holo* funnel, as in the latter, the well of the closed state is expected to be the deeper one (Fig. S8C) (ii) In the case of proteins exhibiting enzymatic functions, the module has to deepen and smoothen the unique closed state well of the *apo* and *holo* funnels, since extreme rigidity is required for catalytic function (Fig. S8D). (iii) The N-terminal module of proteins destined to become transcription factors, by bending-unbending transitions of DNA, is an entire domain that induces oligomerization and tethers the *CC* on DNA, as there is a need to probe a larger conformational space (Fig. S8E). The *CC* protomers have multiple iso-energetic wells corresponding to the different positionings of the N-terminal module with respect to the *CC* (Fig. S8E). Oligomerization and binding to DNA and to their ligands deepens specific wells.

Clearly, the structural promiscuity achieved by the modularity introduced in this study, expands the protein evolvability theory and establish the notion that in order to comprehend protein evolution, it is essential to decode the energetic funnel of structural homologues. Structural elements or even domains added alike Lego bricks to a structural-core, being the Lego Board, trigger distinct evolutionary trajectories.

### Evolution of ligand binding mechanisms

Although extensive structural and biochemical characterization of periplasmic solute binding proteins has improved our understanding of the ligand binding mechanism (lock-and-key, induced-fit and conformational selection) by which various *CCPs* bind their substrates, the evolutionary origin itself is still unknown. Based on our previous work, an induced-fit binding mechanism (or Venus-fly trap^98^) seems the most likely solution for *CCPs* from classes B,C,D,F,G^81^, while class A and E use an apparent lock-and-key mechanism as shown here. Additional insight from this work is that structural embellishments of the *CC* represent likely the evolutionary mechanism by which the large amplitude structural dynamics that underpin how an induced-fit (Venus-Fly trap) emerged. This is most clearly illustrated by the class B proteins, SBD2 in particular (Fig. 2), in which there is a single C-tail helix mediating the only intra-protein interactions that stabilize the open conformation. The new inter-domain interaction would have expanded the conformational space (since the *CC*-protein domain of CynR in class A use a lock-and-key binding mechanism), increasing the conformational diversity of the substrate-free molecular ensemble, and creating conformers able to accommodate non-native substrates. In line with the Tawfik and Tokuriki model, these likely low-abundance structural sub-states may have been selected and enriched leading to the evolution of extant substrate-specific class B proteins. For class C proteins, a similar evolutionary trajectory seems likely. Class G proteins such as MalE, the largest of the proteins we examined, have the additional twist that structural embellishments within the type II PBP structural core, and not only at the C-terminus, may be involved in creating structural dynamics. This raises the possibility that in some class G proteins, the dynamics may also derive from the internal embellishments, freeing the C-tails to take on additional functional roles unrelated to dynamics. A more detailed study of other class G proteins is needed to explore this further. And finally, our results and observation on class E proteins suggest that C-terminal structural embellishments do not always function to enable Tier-0 dynamics. To our knowledge, this is the first proposal of a molecular mechanism for the evolution of the structural dynamics underlying a ligand-binding mechanism.

## Acknowledgments

This work was financed by an NWO Veni grant (722.012.012 to G.G.), an ERC Starting Grant (No. 638536 – SM-IMPORT to T.C.), Deutsche Forschungsgemeinschaft within GRK2062 (project C03 to T.C.), SFB863 (project A13 to T.C.), FWO CARBS (#G0C6814N to A.E. and G.G.). G.G. also acknowledges an EMBO long-term fellowship (ALF 47-2012 to G.G.) and financial support by the Zernike Institute for Advanced Materials and by the Rega foundation. Y.A.M. was supported by the Indonesia Endowment Fund for Education (Lembaga Pengelola Dana Pendidikan Republik Indonesia-LPDP RI PhD scholarship). N.Z. acknowledges an Alexander von Humboldt postdoctoral fellowship. N.E. acknowledges an MSCA SoE FWO fellowship (195872). R.X. was supported by the Chinese Scholarship Council grant. T.C. was supported by the German Academic Exchange Service (DAAD), Center of Nanoscience Munich (CeNS), LMUexcellent and the Center for integrated Protein Science Munich (CiPSM). We thank Eitan Lerner for useful comments and critical reading of the manuscript, Monique Wiertsema for help with smFRET experiments and Florence Husada for support with ITC experiments.

## Author Contributions

GG., M.d.B., Y.A.M, and T.C. conceived and designed research. G.G. and T.C. supervised the project. G.G., K.T., Y.S., and R.X., designed and performed molecular biology. G.G., Y.A.M., R.X., D.A.G., Y.S. and N.E. prepared protein samples. G.G., Y.A.M., M.d.B., N.Z., Y.S. and N.E. conducted single-molecule experiments. A.T. designed and conducted HDX-MS experiments. S.K. designed and conducted MALS and intrinsic tryptophan fluorescence experiments. Y.A.M., M.d.B., R.X. conducted ITC experiments. M.d.B., G.G. and Y.A.M. analysed single-molecule data. G.G. and Y.A.M. conducted structural and evolutionary analysis of proteins and established the structural classification. N.E. and Y.A.M. classified ligands. G.G., Y.A.M., M.d.B., D.A.G., and T.C. interpreted data and wrote the manuscript. G.G. and Y.A.M. prepared figures and artwork. All authors discussed the results and contributed to the final version of the manuscript, which was approved for submission by all authors.

## Author Information

The authors declare no competing financial interest. Correspondence and requests for material should be addressed to G.G. and T.C.

## Materials and Methods

### Structural analysis & Phylogenetic trees

Proteins harbouring the *cherry-core* (*<u>CC</u>*) structure were identified using the “Structure similarity search function” of the PDB web server.^116,117^ Protein structures were aligned using the PDBeFold server.^118^ Additionally, protein structures were downloaded and superimposed using Pymol (Molecular Graphics System, Version 1.6 Schrödinger, LLC) in order to verify and correct manually the results obtained via the PDBeFold server.

The structure-based phylogenetic trees as presented in SuppFile. 2, 6 and summarized in Fig. 1C were created for proteins with known structures using the “all against all structure comparison” feature of the DALI server.^119^ In the summarized sequence- and structure-based phylogenetic trees as presented in Fig. 1C and Fig. S1; the branch of each class has an average length of the branches of all proteins belonging to the very same class obtained from trees presented in SuppFile. 2, 6.

Sequence-based phylogenetic trees as presented in SupFile. 1 and summarized in Figure S1 were computed using the MultiSeq software^120^ embedded within the <u>V</u>isual <u>M</u>olecular <u>D</u>ynamics (VMD 1.9.2) software package.^121^ Briefly, these trees were based on the <u>p</u>ercent of sequence <u>id</u>entity (PID) of polypeptide chains after the structure-based sequence alignment. Given the fact that the *CC* proteins (*CCPs*) share little amount of sequence conservation, MultiSeq was used to optimize the sequence alignment based on the structural alignment template. The alignments of MultiSeq have all been manually inspected.

The sequence-based extended phylogenetic tree as presented in SuppFile. 1 and summarized in Fig. S1 was made using sequence information of proteins with unknown structural information. We used the 53 *CCPs* with known structures to retrieve homologous protein sequences by inspecting the UniProtKB/Swiss-Prot (swissprot) database with Protein Blast.^122^ Our cut-off for sequence coverage was 70%, which yielded ~600 proteins that allowed to construct an extended sequence-based phylogenetic tree. In all analyses server’s default settings were used unless specified.

### Conformational states of the *cherry-core*

Domain movements were detected via the DynDom protein domain motion analysis server^123,124^ for all the proteins that have high resolution structural informational of both structural states available. Interactions between the C-tail and the domains were analysed by submitting the PDB ID codes corresponding to *apo* and *holo* states (Table. S2) in the <u>P</u>rotein <u>I</u>nteractions <u>C</u>alculator (PIC) webserver.^125^ Similarly, the interactions between the N-terminal dimerization helix and the connecting-loop to the *CC* of class A proteins were retrieved and presented in SuppFile. 7. Standard cut-off distances for all interactions were used. The interactions were also manually inspected using Pymol (Molecular Graphics System, Version 1.6 Schrödinger, LLC). Cartoon protein representations, superimposition of structures and distance calculations between residues were accomplished using Pymol (Molecular Graphics System, Version 1.6 Schrödinger, LLC).

### Gene isolation, protein expression and purification

His_10_SBD2 (Class B), OpuACHis_6_ (Class C), MalEHis_6_ (Class G) were expressed and purified as previously described.^126–128^ *CC* of *cyn*R (Class A) gene (UniProtKB-P27111) was isolated from the genome of Escherichia coli DH5α (F– endA1 glnV44 thi-1 recA1 relA1 gyrA96 deoR nupG purB20 φ80dlacZΔM15 Δ(lacZYA-argF)U169, hsdR17(rK–mK+), λ−). The primers were designed to exclude the HTH lysR-type DNA binding domain (1-58 amino acid); thus to include DNA sequence coding the protein chain 59-299 amino acid. Primers introduced *Nde*I and *BamH*I restriction sites, and the gene product was sub-cloned into the pET16 vector (Novagen, EMD Millipore). For (full length CynR, the same procedure was adopted by including amino acids 1-58. CmpA (class E) gene (UniProtKB-Q55460) was isolated from the genome of *Synechocystis sp*. (*strain PCC 6803*). The primers were designed to exclude signal peptide (1-35 amino acid); thus, to include DNA sequence coding the protein chain 36-452 amino acid.

Intergenic region of CynR operon was isolated from the genome of Escherichia coli BL21-DE3 (F– ompT gal dcm lon hsdSB(rB–mB−) λ(DE3 [lacI lacUV5-T7p07 ind1 sam7 nin5]) [malB+]K-12(λS)) using a set of primers, i.e., the forward primers - X379 (5’-CCA TGT TCA GCC ACG GCA AGA AAA TAA TTG ATA TG-3’) and the reverse – X380 (5’-CTG GAA TTT AAG GAA TCC ATC AAT AAT CTC TTT CAC CG-3’). The amplification of the intergenic region results in the DNA fragment that includes 191 bp, which subsequently was used as the binding partner of CynR for further experiments.

Protein derivatives having the cysteine mutations or the point mutations indicated into brackets throughout the manuscript were constructed using QuickChange mutagenesis^129^ and Megaprimer PCR mutagenesis^130^ protocols.

His_10_CynR (*CC*) was over-expressed in BL21 DE3 cells (F– ompT gal dcm lon hsdSB(rB–mB−) λ(DE3 [lacI lacUV5-T7p07 ind1 sam7 nin5]) [malB+]K-12(λS)). Cells harboring plasmid expressing the protein were grown in Terrific Broth medium (30°C; OD_600 nm_=0.5) and protein overexpression was induced by IPTG (0.10 mM; growth at 16°C for 15 hours).

His_10_CmpA was over-expressed in BL21 pLysS DE3 cells (F–, *omp*T, *hsd*S_B_ (r_B_–, m_B_−), *dcm*, *gal*, λ(DE3), pLysS, Cm^r^). Cells harboring plasmid expressing the protein were grown (30°C; OD_600 nm_=0.5) and protein overexpression (growth at 16°C for 15 hours) was induced by IPTG (0.1 mM). For all proteins growth was accomplished in Luria Bertani medium, except for CynR in Terrific Broth.

Proteins were purified as follows: after lysis by cell disruptor (30.000 psi; 2 rounds), soluble material (50.000 g; 30 min; 4 °C; Sorval) was loaded (50 mM Tris-HCl, pH = 8; 1 M KCl, 10% glycerol; 10 mM Imidazole; 1 mM DTT; 2 mM PMSF) on a Ni-NTA resin (Qiagen). Bound proteins were washed (50 mM Tris-HCl, pH = 8; 50 mM KCl, 10% glycerol; 10 mM Imidazole; 1 mM DTT; and 50 mM Tris-HCl, pH=8; 1 M KCl, 10% glycerol; 30 mM Imidazole; 1 mM DTT sequentially) and then eluted (50 mM Tris-HCl, pH=8; 50 mM KCl, 10% glycerol; 300 mM Imidazole; 1 mM DTT). Protein fractions were pooled (supplemented with 5 mM EDTA; 50 mM DTT), concentrated (Vivacell-Sartorious; Amicon-Millipore), dialyzed (50 mM Tris-HCl, pH=8; 1 M KCl, 50% glycerol; 10 mM DTT), aliquoted and stored at −80°C.

For untagged CynR (Full length): after lysis by cell disruptor (30.000 psi; 2 rounds), soluble material (50.000 g; 30 min; 4 °C; Sorval) was loaded (50 mM Tris-HCl, pH = 8.3; 5mM NaCl, 3 mM EDTA; 1 mM DTT; 2 mM PMSF) on a Q sepharose fast flow (GE Healthcare) and incubated over-night. Immobilized proteins were washed (50 mM Tris-HCl, pH 8.3, 135 mM NaCl, 3mM EDTA, 1mM DTT) and eluted (50 mM Tris-HCl, pH 8.3, 250 mM NaCl, 3mM EDTA, 1mM DTT). Protein fractions were pooled and concentrated (10.000 MWCO Amicon; Merck-Millipore) up to 5 mL and analyzed on a HiLoad 26/60 Superdex 200 (GE) (equilibrated with 50 mM Tris-HCl, pH 8.3, 5 mM NaCl, 0,1mM EDTA). Fraction were analyzed on SDS-PAGE and those containing pure CynR were collected and re-subjected to Size-exclusion chromatography to increase purity (>90%). Fractions were collected, dialyzed (50 mM Tris-HCl, pH 8.0, 200 mM NaCl, 50% glycerol, 0,5mM EDTA and 1 mM DTT), aliquoted and stored at −20°C.

### Protein labelling

Protein labelling was accomplished as described previously.^126^ In brief, cysteine positions were chosen based on the open and closed x-ray crystal structures of CynR (2HXR, 3HFU), OpuAC (3L6G, 3L6H), SBD2 (4KR5, 4KQP), CmpA (2I49, 2I4C) and MalE (1OMP, 1ANF); Table. S5. Stochastic labelling was performed with the maleimide derivative of dyes Cy3B (GE Healthcare) and ATTO647N (ATTO-TEC) for OpuAC and SBD2. CmpA, CC of CynR and MalE were labelled with Alexa555 and Alexa647 (ThermoFischer). Histidine tagged proteins were immobilized on a Ni^2+^-sepharose resin (GE Healthcare) to remove the 1 mM DTT (Dithiothreitol) used to keep all cysteine residues in a reduced state. The resin was incubated 2-8 hours at 4°C with 50 nmol of each fluorophore dissolved in the appropriate buffer (50 mM Tris-HCl, pH=7.4; 50 mM KCl for CynR (CC), CmpA and MAlE; 50 mM KPi, pH=7.4; 50 mM KCl for SBD2 and OpuAC) and subsequently washed to remove the majority of unbound fluorophores. The labelled protein was further analysed by size-exclusion chromatography (Superdex 200, GE Healthcare) to enrich the double-labelled fraction and remove potential aggregation material^131^. For all proteins, labelling efficiency was higher than 80%.

### Isothermal titration calorimetry (ITC)

Purified CynR was dialyzed overnight against 50 mM Tris-HCl, pH=7.4; 50 mM KCl. ITC experiments were carried out in a Nano ITC Low Volume (TA Instruments). The sodium azide stock solution (250 μM) was prepared in the dialysis buffer and was stepwise injected (0.5 μl) into the reaction cell containing 12 μM CynR. All experiments were carried out at 298 K with a mixing rate of 250 r. p. m. Data were analysed with a single-binding site equation, provided by the NanoAnalyse software (TA Instruments).

Purified CmpA was dialyzed overnight against 1X PBS Buffer that has been supplemented with 50uM K_2_CO_3_ and 1uM EDTA. ITC experiments of CmpA were performed using MicroCal iTC200 (Malvern). The Calcium solution (500uM) was prepared from CaCl_2_ powder, which subsequently diluted in the dialysis buffer and was injected (2 μl) into the reaction cell containing 40 μM of CmpA. All experiments were carried out at 297 K with a mixing rate of 750 rpm. Data were analysed with a single-binding site equation, provided by the provided MicroCal Analysis software (Malvern).

### Solution-based smFRET and ALEX

ALEX experiments were carried out at 25-50 pM of double-labelled protein in the appropriate buffer (50 mM Tris-HCl, pH=7.4; 50 mM KCl for CynR, CmpA and MalE; 50 mM KPi, pH=7.4; 50 mM KCl for SBD2 and OpuAC) supplemented with additional reagents as stated in the text. The experiments were performed using a home-built confocal microscope similar to the setup described before^126^ with minor modifications as outlined below. Briefly, two laser-diodes (Coherent Obis) with emission wavelength of 532 and 637 nm were modulated in periods of 50 µs and used for confocal excitation. Alternation between both excitation wavelengths was achieved by direct modulation of the two laser. The beam of both lasers was coupled into a single-mode fiber (PM-S405-XP, Thorlabs) and collimated (MB06, Q-Optics/Linos) before entering a water immersion objective (60X, NA 1.2, UPlanSAPO 60XO, Olympus). The excitation spot was focused 20 µm above the interface of glass and water solution. Typical average laser powers were 30 μW at 532 nm (~30 kW/cm^2^) and 15 μW at 637 nm (~15 kW/cm^2^). Excitation and emission light were separated by a dichroic beam splitter (zt532/642rpc, AHF Analysentechnik), mounted in an inverse microscope body (IX71, Olympus). Emitted light was focused onto a 50 µm pinhole and spectrally separated (640DCXR, AHF Analysentechnik) onto two APDs (τ-spad, <50 dark-counts/s, Picoquant) with appropriate spectral filtering (donor channel: HC582/75; acceptor channel: Edge Basic 647LP; both AHF Analysentechnik). Photon arrival times were registered by an NI-Card (PXI-6602, National Instruments). A dual-colour burst search^132^ using parameters M = 15, T = 500 μs and L = 25 was applied to identify bursts. Additional thresholding was applied to remove spurious changes in fluorescence intensity and selected for intense single-molecule bursts (total photons per burst > 150 unless otherwise mentioned). Binning the detected bursts into a 2D apparent FRET/S histogram (61 × 61 bins unless otherwise mentioned) allowed the selection of the donor and acceptor labelled molecules. ^133^ The selected apparent FRET histograms (61 × 1 bins unless otherwise mentioned) were fitted using a Gaussian function.

Single-molecule bursts from donor-only labelled CynR and MalE proteins were obtained by selecting the donor-only subpopulation (S > 0.9) from the 2D apparent FRET/S histogram. The total photon count per burst were normalised by its respective duration to obtain the photon count rate.

### Scanning confocal microscopy

Confocal scanning experiments were performed using the same homebuilt confocal microscope as described before.^126^ Surface scanning was performed using a piezo stage with 100×100×20 μm scanning range in XYZ, respectively (P-517-3CD with E-725.3CDA, Physik Instrumente). The detector signal was registered using a Hydra Harp 400 picosecond event timer and a module for time-correlated single photon counting (both Picoquant). Data were recorded with constant 532-nm excitation at an intensity of 0.5 μW (~125 W/cm^2^) for SBD2 and MalE, but 1.5 μW (~400 W/cm^2^) for OpuAC. Data acquisition was done with home-written Labview software (National Instruments). A flow-cell arrangement was used for studies of surface-immobilized OpuAC and SBD2; whereas MalE was studied on standard functionalized cover-slides. Flow-cells were assembled using established protocols described before.^126,134^ All experiments of OpuAC and SBD2 were carried out in a degassed buffer (50 mM KPi, pH=7.4; 50 mM KCl) under oxygen-free conditions obtained utilizing an oxygen-scavenging system supplemented with 10 mM of aged Trolox as a photostabilizer.^135^ For MalE, experiments were carried out in buffer (50 mM Tris-HCl, pH=7.4; 50 mM KCl) supplemented with 1 mM Trolox.^135^

Time-traces were analysed by integrating the detected red and green photon streams in time-bins of 1.5 ms (OpuAC) or 5 ms (MalE and SBD2). Only traces lasting longer than 50 time-bins having on average more than 10 photons per time-bin were used for further analysis. The apparent FRET per time-bin was calculated by dividing the red photons by the total number of photons per time-bin. The most probable state-trajectory was identified by Hidden Markov Modeling (HMM)^136^. For this an implementation of HMM was programmed in Matlab (MathWorks), based on previous studies.^136^ We assume that the FRET time-trace (the observation sequence) can be considered as a HMM with only two states having a one-dimensional Gaussian-output distribution. The aim of a HMM is to infer both the states and state-transition probabilities given the observation sequence. The Gaussian output-distribution of a state *i* (*i* =1, 2) is completely defined by two parameters: the average and the variance. The HMM algorithm finds all parameters λ, given only the observation sequence, that maximizes the likelihood function.^136^ This was done with the forward-backward algorithm.^137^ Care was taken to avoid floating point underflow and was done as described.^136^ The most probable state-trajectory is then found from λ by using the Viterbi algorithm.^138^ The time spend in each state is interfered from the most probable state-trajectory and is summarized into a histogram for all traces under the same condition. The histogram was fitted with a single exponential decay to obtain the closing rate and closed-state lifetime.

### Hydrogen/Deuterium exchange Mass Spectrometry (HDX-MS)

#### Isotope labeling

MalE and derivative MalE (M321K) were incubated for 30 min at 25°C [5 μl of 28.25 μM protein stock in 50 mM Tris-HCl pH 7.4, 50 mM KCl to ensure saturation and to thermally equilibrate the samples. At 30 min of incubation, MalE was isotopically labeled with 93.75% v/v final D content at 25°C, by addition of 75 μl deuterated buffer [50 mM Tris-DCl pD 7.4, 50 mM KCl in D_2_O (D_2_O, 99.9% atom D; Euriso-top)] for 10, 100, 1000, 10000 and 100000 sec. pD refers to the corrected value for the isotope effect. The HDX reaction was quenched at the defined time intervals by instant acidification [pD 2.5; formic acid; (Ultra-pure) from Merck KGaA]. The pre-chilled quenching solution contained urea (Urea-d4; Sigma) to a final concentration of 1.6 M at quenching to increase the peptide coverage by mild denaturation. 50 pmoles of protein were injected for analysis.

The CC of CynR (92-298) and the tetrameric DNA-free CynR (1-298) (isolated by SEC) were concentrated to 28.6 uM and supplemented with 5 mM DTT. CynR (4 μl; 28.6 μM) was incubated at 25°C for 30 min in the presence or absence of NaN_3_, by addition of 1 μl of 0.5 M NaN_3_ diluted in 50 mM Tris-HCl pH 7.4, 50 mM KCl. For the apo states, 1 μl of buffer was added instead. Deuterated buffer (95 μl; as for MalE) was added for 10, 100, (1000/ 2000 for the CC and CynR respectively), 10000 and 100000 (for the CC) sec at 25°C, resulting in 95% v/v D_2_O, 1.14 μM FL CynR, 5 mM NaN_3_. The isotope labeling reactions were quenched by acidification (pD_corrected_ 2.5) with pre-chilled quenching solution containing urea (Urea-d4; Sigma) to a final urea concentration of 1.52 M (0.73 μM protein). The samples were instantly frozen in liquid nitrogen and analyzed by instant thawing within two days of preparation. 36 pmoles of protein were injected for analysis.

For the analysis of the DNA-bound CynR, a different approach was followed to minimize the signal suppression during the mass spectrometric analysis induced by the addition of DNA. Specifically, FL CynR (2 μl; 32.85 μM protein; 5 mM DTT) was supplemented with 1.2 molar excess of DNA in H_2_O (1.8 μl of 43.99 μM) and incubated at 25°C for 5 min prior to addition of 0.6 μl of 0.5 M NaN_3_ (in H_2_O) for the liganded state, or with 0.6 μl of nanopure H_2_O for the apo state. The dilution of Tris and KCl concentrations by addition of H_2_O at this step was corrected by increased their concentrations in the protein stock. The samples were incubated for an additional 25 min at 25°C prior to addition of deuterated buffer (55.6 μl; as above) for 10, 1000, 10000 sec at 25°C, resulting in 92.7% v/v D_2_O, 1.1 μM CynR, 1.3 μM DNA, 5 mM NaN_3_. The isotope labeling reactions were quenched by acidification (pD_corrected_ 2.5) with 4.2 excess of pre-chilled quenching solution containing deuterated urea to a final concentration of 1.52 M (250 μl total volume of quenched reaction; 0.26μM CynR; 0.32μM DNA) and loaded on centrifugal concentrators (Vivaspin 500; 10,000 MWCO PES, Sartorius). The quenched samples were concentrated to 60 μl by centrifugation at 20,000xg for 2 min (Sigma 1-16K). Due to the linear nature of DNA, it is not retained in the centrifugal membrane and therefore only the protein is concentrated (1.1 μM CynR; 0.32μM DNA). The retained solution in the centrifugal apparatus was transferred to a pre-chilled tube and was frozen in liquid nitrogen.

#### Online proteolysis-LC-MS analysis

The quenched samples were injected into a nanoACQUITY UPLC System with HDX technology (Waters, UK), thermostated at the digestion and LC separation chambers at 20°C and 0.8°C (2°C for CynR) respectively. Proteolytic digestion (Enzymate BEH pepsin column, Waters) and peptide trapping/desalting (ACQUITY UPLC R BEH C18 VanGuard pre-column; 130 Å, 1.7 μm, 2.1 × 5 mm; Waters) were performed with 0.23% formic acid in H_2_O (Solvent A) at 100 μl/min for 3 min, online with peptic peptide separation (ACQUITY UPLC R BEH C18 analytical column; 130 Å, 1.7 μm, 1 x 100 mm; Waters) at 40 μl/min using a 12 min (8 min for CynR) linear gradient from 5% to 50% Solvent B [ACN (Optima LC/MS grade; Fischer Scientific), 0.23% formic acid]. The eluate was analyzed online on a Synapt G2 ESI-Q-TOF instrument (Waters, UK) with a MassLynX interface (version 4.1 SCN870; Waters) for data collection. The source/TOF conditions were set as: resolution mode, capillary voltage 3.0 kV, sampling cone voltage 20 V, extraction cone voltage 3.6 V, source temperature 80°C, desolvation gas flow 500 L/h at 150°C. The deuterated samples were analyzed in MS acquisition mode (300-2000 Da range), while for peptide identification non-deuterated samples (treated as above but in protiated buffers) were analyzed in MS^E^ acquisition mode over the m/z range 100-2,000 Da, using a collision energy ramp from 10 to 30 V. Leucine Enkephalin (2 ng/μl in 50% ACN, 0.1% formic acid; 5 μl/min) was co-infused in both acquisition modes for accurate mass measurements (reference mas: m/z 556.2771).

#### Data analysis

For peptide identification, MS^E^ data were processed on the ProteinLynx Global Server (PLGS v3.0.1, Waters, UK), using a user-defined database containing MalE, MalE (M321K), the CC CynR and CynR sequences under the following criteria: digestion enzyme, non-specific; false discovery rate, 4%, minimum fragment ion matches/peptide and /protein, 3 and 7; minimum peptide matches/protein, 1; low and elevated energy thresholds, 150 and 25 counts; intensity threshold: 500 counts, reference mass correction window, 0.25 Da at 556.2771 Da/e. The identified peptides from two independent MS^E^ raw files were further filtered on the DynamX software (version 3.0, Waters) for DynamX score> 7.5 (7 for CynR), maximum MH+ error of 5 ppm and minimum products/amino acid of 0.2. Only robustly identified peptides in both replicates were further processed, resulting in 94.3, 93.8%, 93.4% and 88.3% sequence coverage for MalE, MalE (M321K), the CC CynR and full-length CynR respectively. Deuterium uptake was determined using the DynamX software. For the comparison of the WT and MalE (M321K) at the mutated region, the peptide containing residues 321-337 was compared only at the level of D-uptake relative to the full deuteration of each state, after complete analysis of each protein with its individual controls, and not at the level of absolute D-uptake. For this, MalE (M321K) was analyzed using the peptide list created from the MalE (M321K) sequence, the MalE (M321K) non-deuterated and full deuterated controls (Table F5B). Additionally, for the comparison of MalE (M321K) at the apo state, statistical analysis was performed both on the data analyzed using the WT and MalE (M321K) sequence in order to include the mutated region (Tables F5B, F5C, F5D).

#### Statistical analysis

All HDX reactions were performed in triplicates, unless otherwise indicated (Table F5A, F4A, F4B, F4C). Statistical analysis of the significance of differences between the apo and liganded states, and between the WT and MalE (M321K) mutated sequence in the case of MalE, was achieved using a modified approach of Bennett *et al*., ^139^, described in Tsirigotaki *et al*., ^140,141^ Briefly, two-tail paired t-tests, comparing the mean uptake, as absolute deuterium uptake values, of the two states for each peptide, were performed using R language and the significance threshold was set to 99% confidence (1-p≥0.99). An additional threshold was set at ±4SD of average pooled standard deviations of both states. Finally, a third criterion was introduced to exclude false positives due to high SD outliers that are averaged out in the pooled SDs: The difference between the two states for each peptide must exceed twice the sum of SDs of the two states for the given peptide. Only differences that fulfill all three criteria were considered as statistically significant. Visualization of the statistical analysis in scatter plots was achieved by R language (Table F5D, F4A, F4B and F4C). For optimal realization of the extent of the ΔD-uptake regardless of the length of the identified peptides, the ΔD-uptake of each peptide was expressed relative to the experimentally determined full deuteration control (normalized ΔD-uptake). For higher robustness, only statistically significant differences that exceed the absolute value of 10% normalized ΔD-uptake are discussed. Detailed data sets including smaller, statistically significant differences are given in Tables F5C (for WT and MalE (M321K) apo states), F5E (for WT MalE apo and holo states) F4A (for DNA-bound CynR apo and holo states), F4B (for DNA-free CynR apo and holo states) and F4C (for the CC CynR apo and holo states).

### Bulk Fluorescence measurements

Measurements were carried out in quartz cuvettes (1 ml; Hellma) with a Cary Eclipse fluorimeter supplemented with a four-position cuvette holder and a Peltier temperature controller (Varian). To determine an apparent meting temperature T_m (apparent)_, the changes in intrinsic tryptophan fluorescence emission of CynR (0.3 μM) as a function of increasing temperature (10 - 70 °C; ramping rate 0.8 °C/min; excitation 297 nm/emission 345 nm; slits at 2.5 and 20 nm; data acquisition interval = 0.5 min), in the presence or absence of NaN_3_ (1 mM). All data were collected using Cary Eclipse software (Bio Package; Varian) and analyzed by nonlinear regression using Origin 5.0 (Microcal).

### Homology modelling method

For establishing the three-dimensional structure and the different oligomeric states of CynR, its amino acid sequence has been modelled according to known structures having a substantial degree of sequence similarity that were used as templates^142^. The amino acid sequence of CynR was submitted to the SWISS-MODEL in the Automated Protein Modelling server provided by the Glaxo Smith Kline center (Geneva, Switzerland) using the standard settings of the server^143^. All models were reliable according to server thresholds^144–146^.

## Supporting Information

**Figure S1:**
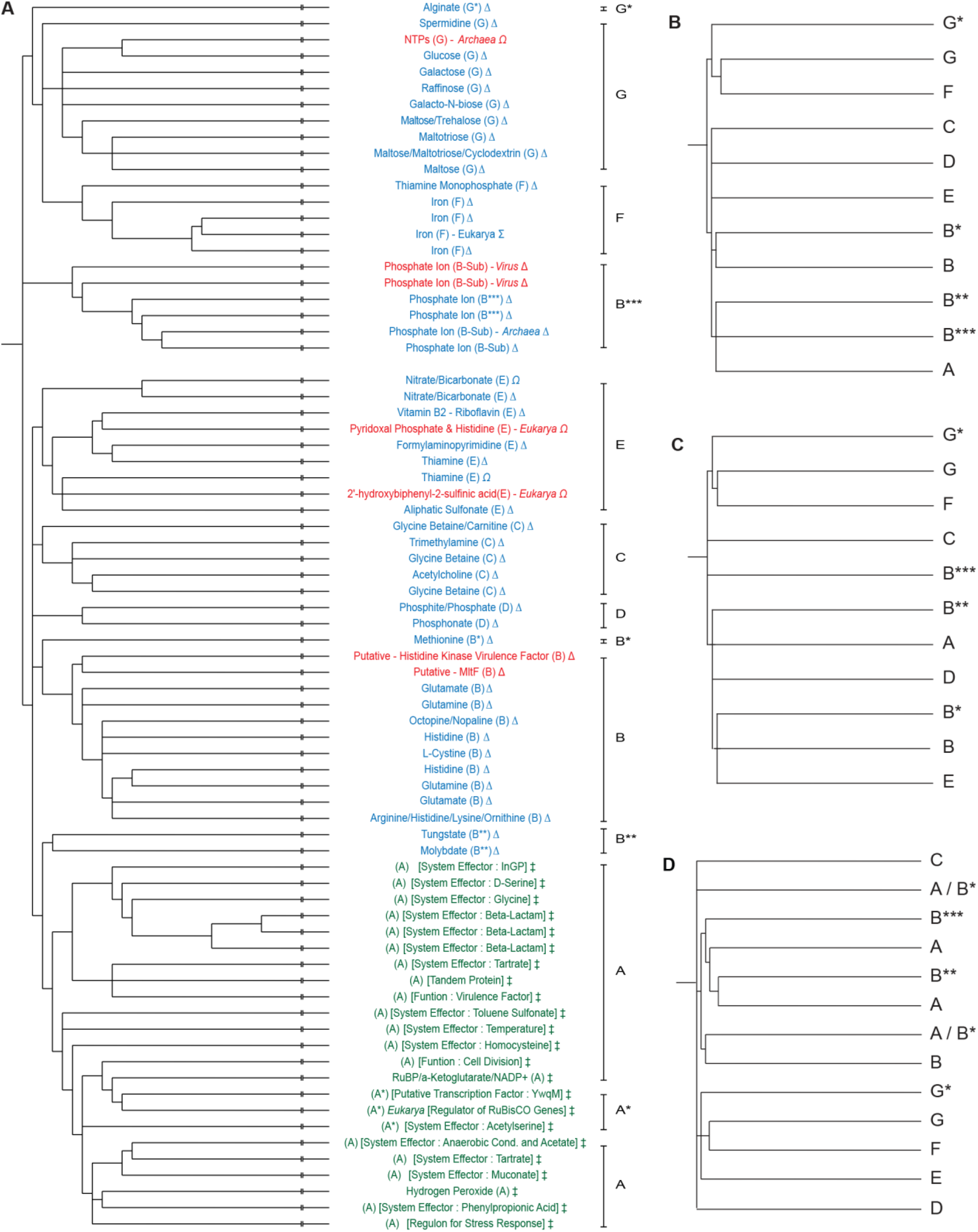
Phylogenetic Trees of *cherry-core* proteins, related to Figure 1. **(A)** Summarized sequenced-based extended phylogenetic tree of the bilobed *cherry-core* proteins (*CCPs*), grouped with respect to their ligands. Complete tree of ~ 600 proteins in shown in SuppFile. 1. Function was obtained from UniProtKB^1^ and is color-coded (Green: transcription factors; Red: enzymes; Blue: substrate binding proteins). Localization is denoted by a symbol (‡: DNA-bound; Δ: extracellular; Ω: cytoplasm; Σ: chloroplast). Ligands or system effectors are indicated and chemical structure is presented in Fig. S3. **(B-D)** Summarized (complete trees in SuppFile. 2) sequenced-based phylogenetic trees of the 53 identified *CCPs* with known structures for the entire polypeptide chain **(B)**, for D1, D2 **(C)**, and for the C-tail only **(D)**. An asterisk (*) marks structural subclasses (see SuppFile. 3). A similar clustering is observed in all above trees and comparable to the structure-based phylogenetic tree (Fig. 1C). This indicates that proteins having a class-specific rigid domain (D1 and D2) have evolved to have the class-specific C-tail. This outcome is in line with the fact that the C-tail–domain interactions stabilize a define structural state (Fig. 2 and Table. S2)

Two major clades are encountered in the structure-based phylogenetic trees (Fig. 1C). The ‘lower clade’, which includes classes F and G, manifest substantial alterations of the *CC*, represented primarily by insertions of secondary structural elements (see also Fig. 1D and Fig. S2). Our secondary structural alignments indicated that the most pronounced alterations in classes F, G involve: the addition of <u>α</u>-helix <u>3</u> (α3), α-helix 4 (α4) and the deletion of β-sheet 5 (β5). On the other hand, the ‘upper clade’ includes *CCPs* with minor alterations/variability of their *CC*. In this study, we selected to analyse representative examples to describe faithfully the phylogenetic relationships: From the lower clade, MalE of Class G has been selected as it fulfils the following criteria: (i) belongs to the class with the most complex C-tail (Fig. 1C, D) of the *CCPs* and (ii) includes some of the most well-studied proteins with abundant structural information available in the Protein Data Bank. OpuAC of Class C has been selected as it belongs to the first diversified branch of the ‘upper clade’. Moreover, SBD2 of Class B has been picked regarding its long evolutionary distance with the previously selected ‘upper clade’ cousin (i.e. OpuAC). CynR (Class A) and CmpA (Class E) have also been analysed experimentally as the only ones with no Tier-0 dynamics (Fig. 2 and Table. S1).

**Figure S2:**
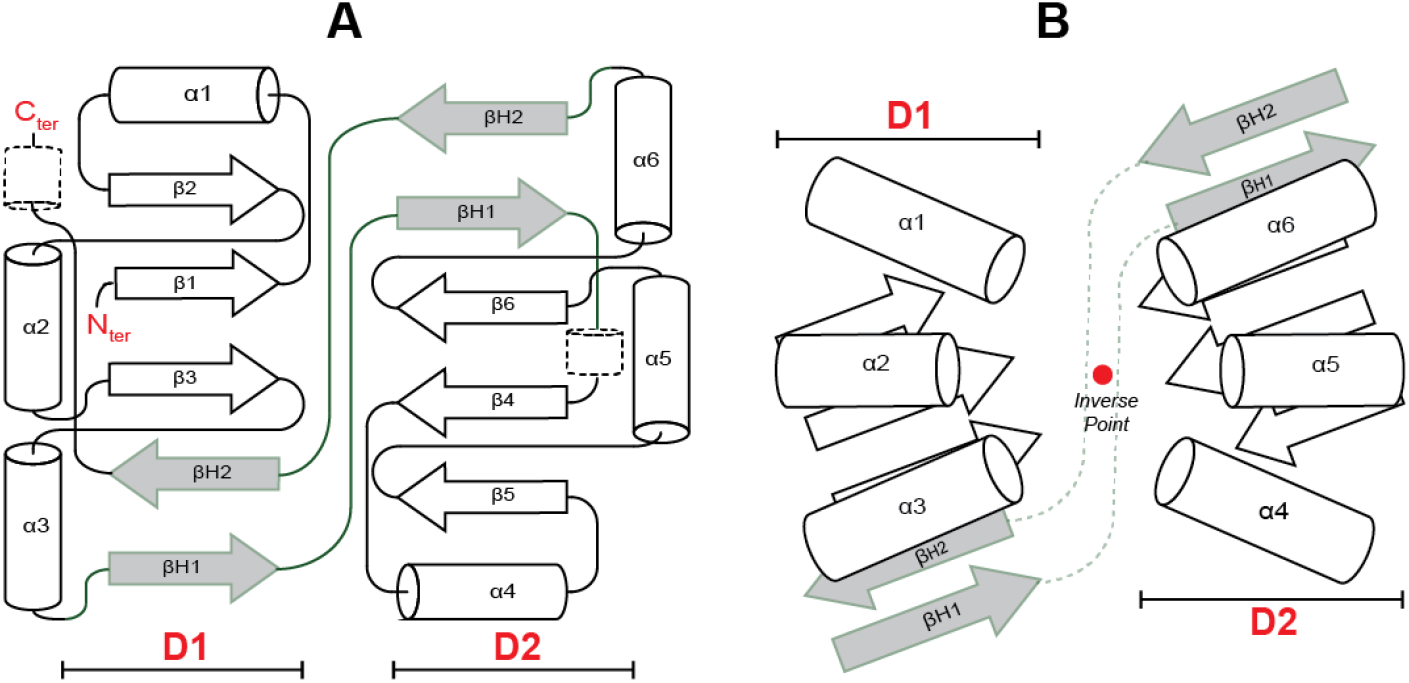
Structural Consensus of *cherry-core* proteins, related to Figure 1B-D. **(A)** Consensus/model derived from secondary structure alignments of the *CC* of *CCPs* (Fig. 1D and SuppFile. 3). **(B)** Schematics of the geometrical consensus for the 3D placement of secondary structure elements originated from the 3D Structural analysis (manually inspected by Pymol). The unique placement of such elements in the 3D space provides a remarkable degree of symmetry of D1 & D2 of the *CCPs* forming the binding site.

**Figure S3.**
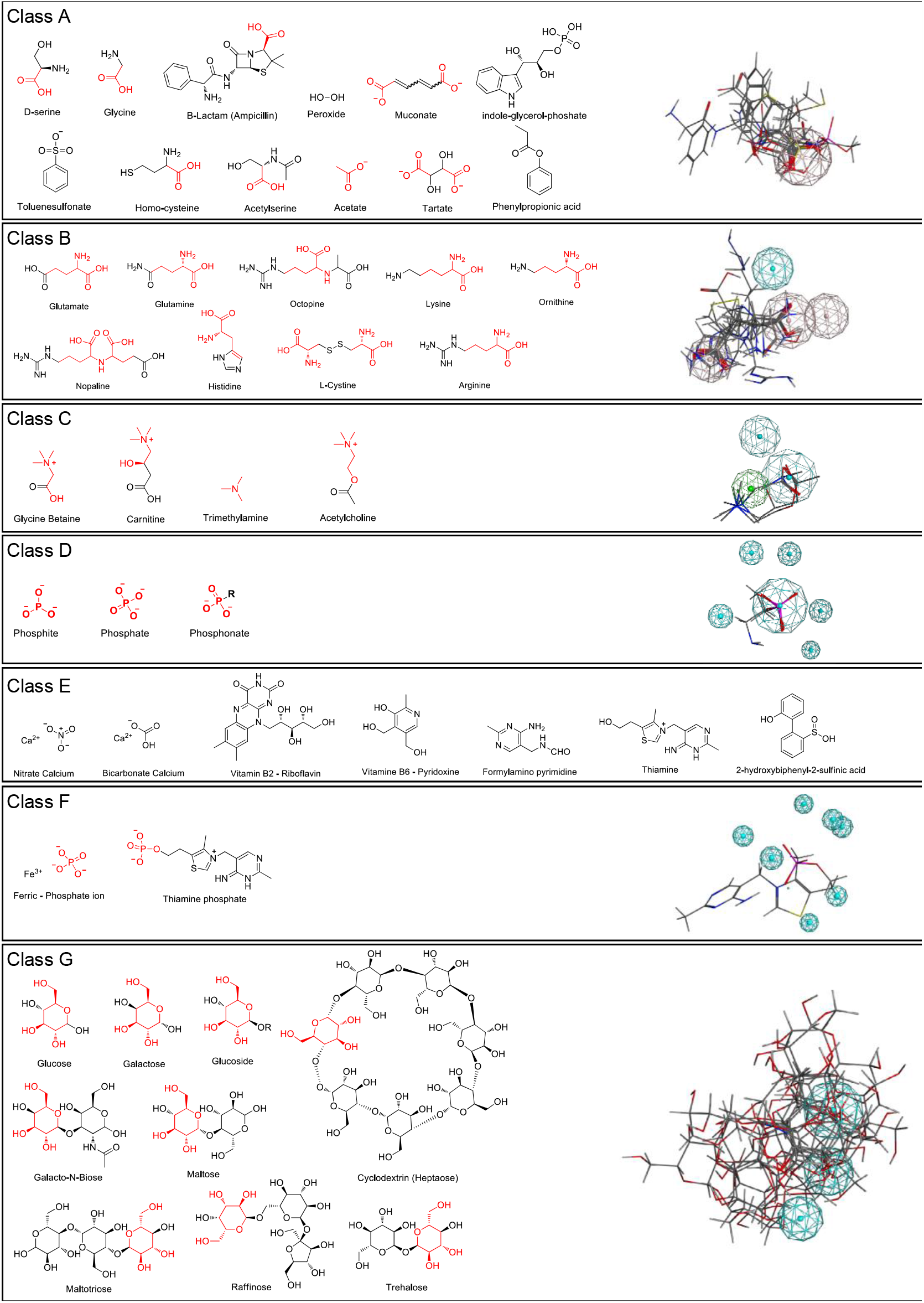
Common Chemical Structure and pharmacophore structure of the *cherry-core* protein ligands, related to Figure 2. The common chemical structure of all identified ligands for each of the different *CCP* classes (A-G) is highlighted with red color. The ligands of class A, but mostly class E do not share a substantial degree of a chemical structure. In addition, pharmacophore analysis, after flexible alignment (by using the MOE software) of the ligands for every class, reveals the common interactions which are represented with brown (hydrogen bond donor-acceptor, HBD-A), blue (hydrogen bond acceptor, HBA) or green (lipophilic) spheres. Specifically, class A shares 1 interaction (HBD-A); class B, 4 interactions (3 HBD-A and 1 HBA); class C, 3 interactions (2 HBA and 1 lipophilic); class D, 6 interactions (HBA); class E, 0 interactions; class F, 7 interactions (HBA) and class G, 3 interactions (HBA).

**Figure S4:**
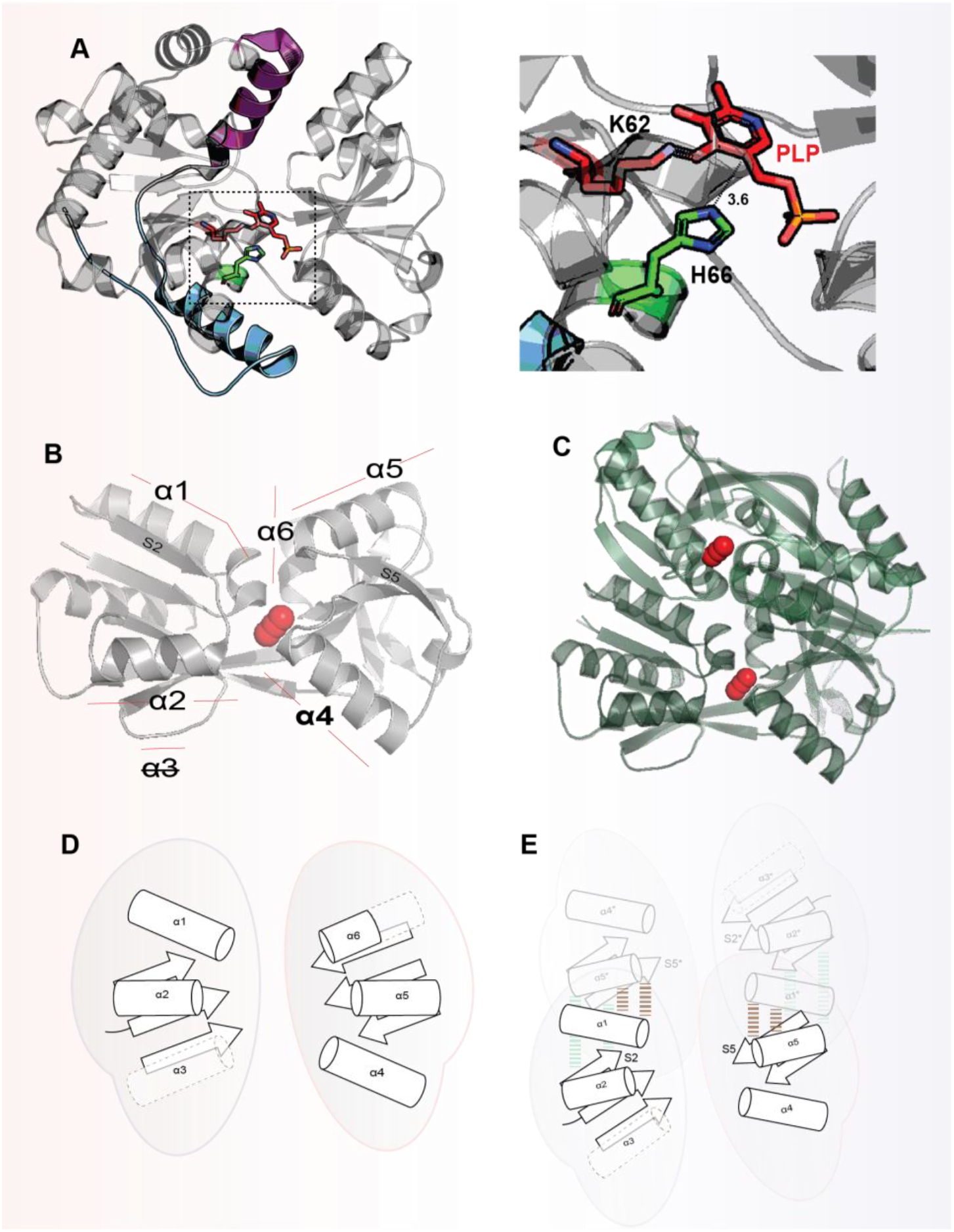
Structural Analysis of the oligomeric transcription factor CynR and of the single-turnover enzyme pyrimidine synthase THI5p, related to Figure 2, 4. **(A)** Crystal structure (PDB:4ESX) of the single-turnover enzyme thiamin pyrimidine synthase^2^ in complex with its ligand pyridoxal phosphate (red sticks). The 2 symmetric helices of the C-tail that stabilize the cleft in the closed structure and render it rigid, are colored as in Fig. 2. The extreme rigidity required is evidenced by the fact that pyridoxal phosphate needs to bind to the cleft of the pyrimidine synthase of *C. albicans* with a very specific geometry (provided also by the covalent interaction with K62, forming a Schiff base) with respect to the side chain of H66 (green sticks). This allows the “remarkable” chemistry to occur, as stated by the authors of the corresponding paper. The reaction consists of the excision of the side chain of H66 to produce thiamine pyrimidine. **(B)** Crystal structure of the *holo* state of the *CC* of CynR (PDB: 3HFU) with azide (indicated with red spheres) in the binding cleft. For clarity, the second protomer present within the assymetric unit, was omitted. **(C)** The *holo* state of the *CC* CynR (PDB: 3HFU, grey color) was superimposed with the *apo CC* CynR (PDB: 2HXR, green color). The dimeric assembly was classified to be biologically relevant by EPPIC. Both the protomer structure, and the dimeric assembly do not undergo significant structural changes and are perfectly super-imposable. **(D)** Schematic representation of the *CC* of CynR. The conserved *CC* helix (Fig. S2) α3 is missing from the *CC* of CynR and all class A proteins (Fig. 1D, SuppFile. 3). **(E)** Schematic representation of the dimeric *CC* of class A proteins. Interacting secondary structural elements are highlighted. α1 makes hydrophobic contacts with S5 and α5, while S2 hydrophobic contacts with α5 and hydrophobic contacts/hydrogen bonds with S5.

**Figure S5.**
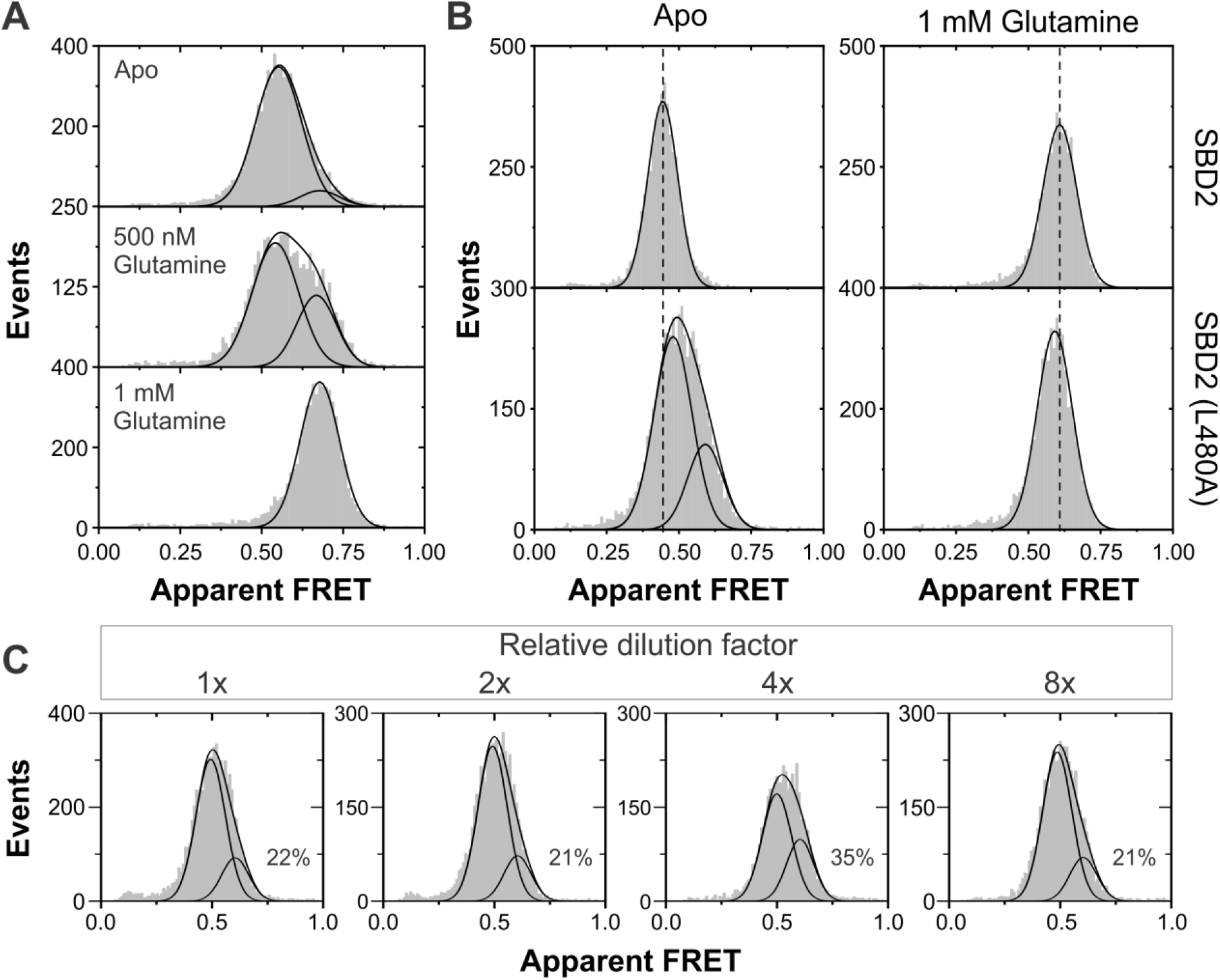
Conformational dynamics of SBD2 and derivatives, related to Figure 5 (A-D). **(A)** Apparent FRET efficiency histogram of freely diffusing SBD2 (L480A) labelled with the donor fluorophore Alexa Fluor 555 and acceptor fluorophore Alexa Fluor 647. The fraction of closed state is similar as found for SBD2 (L480A) labelled with the donor fluorophore Cy3B and the acceptor fluorophore Atto647N, as presented in Fig. 5C. **(B)** The apparent FRET efficiency for both the *apo* (open) and *holo* (closed) state of SBD2 (L480A) is found to differ slightly from SBD2, which would suggest that the energy landscape is altered. **(C)** To exclude that the presence of endogenous ligand is responsible for the fraction closed observed, FRET efficiency histograms were measured under different sample concentrations. By diluting the sample, the ligand concentration would decrease, and hence the fraction of closed observed should decrease if it was due to ligand contamination. These results show no change in fraction closed over almost a 10-fold increase in dilution, indicating that the fraction closed is not caused by endogenous ligand.

**Figure S6.**
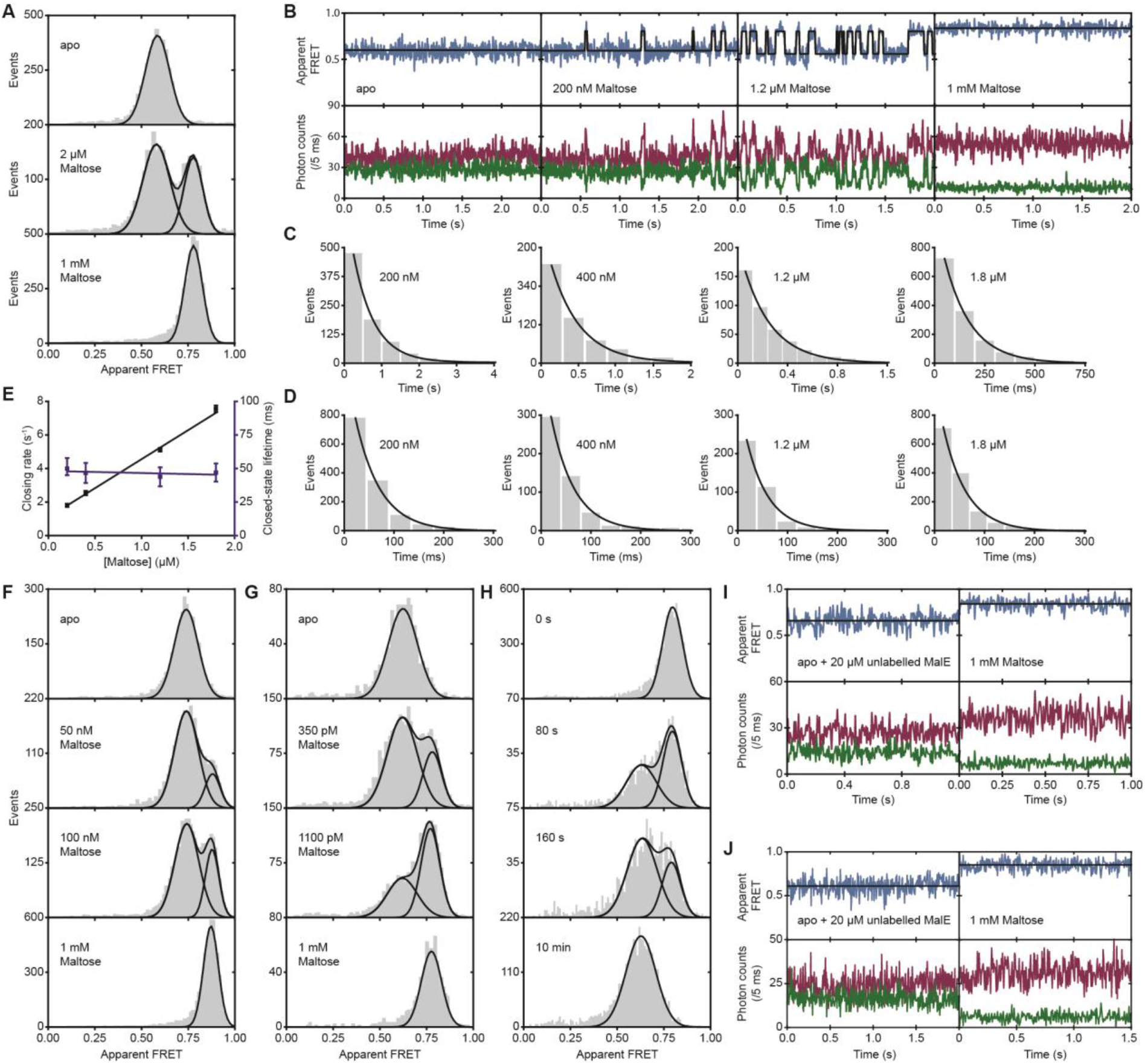
Conformational dynamics of MalE and derivatives, related to Figure 5 E-I. **(A)** Apparent FRET efficiency histogram of MalE, obtained from the solution-based smFRET and ALEX measurements under conditions as indicated. **(B)** Fluorescence trajectories of MalE under different conditions as indicated; donor (green) and acceptor (red) photon counts are binned with 5 ms. The top panel shows calculated apparent FRET efficiency (blue) with the most probable state-trajectory of HMM (black). **(C, D)** Waiting time distribution of the low (**C**) and high FRET state (**D**) as obtained from the most probable state-trajectory of the HMM of all molecules per condition. Grey bars are the binned data and the solid line is the fit. **(E)** Mean closing rate (black) and closed-state lifetime (purple) as function of maltose concentration obtained from the fits in (C) and (D). Error bars indicate the 95% confidence interval of the fit obtained from the fit in (C) and (D). **(F, G)**: Apparent FRET efficiency histogram of MalE (M321A) (**F**) and MalE (M321K) (**G**). **(H)** Time dependence of apparent FRET efficiency histograms of MalE (M321K) in the presence of 10 nM maltose and 20 μM unlabeled MalE (M321K) added to scavenge all maltose. First 10 nM maltose was added to saturate the protein (top panel). Subsequently ~20 µM unlabeled protein was added to scavenge all ligands that are released from the labelled protein. Protein conformation was probed at indicated time points. Grey bars are the data and solid line is the fit. **(I, J)** Fluorescence trajectories of MalE (M321A) (**I**) and MalE (M321K) (**J**) obtained from surface immobilized molecules in the *apo* (~20 μM unlabeled protein) or 1 mM maltose conditions. Fluorescence trajectories of MalE derivatives are plotted similarly to (B).

**Figure S7:**
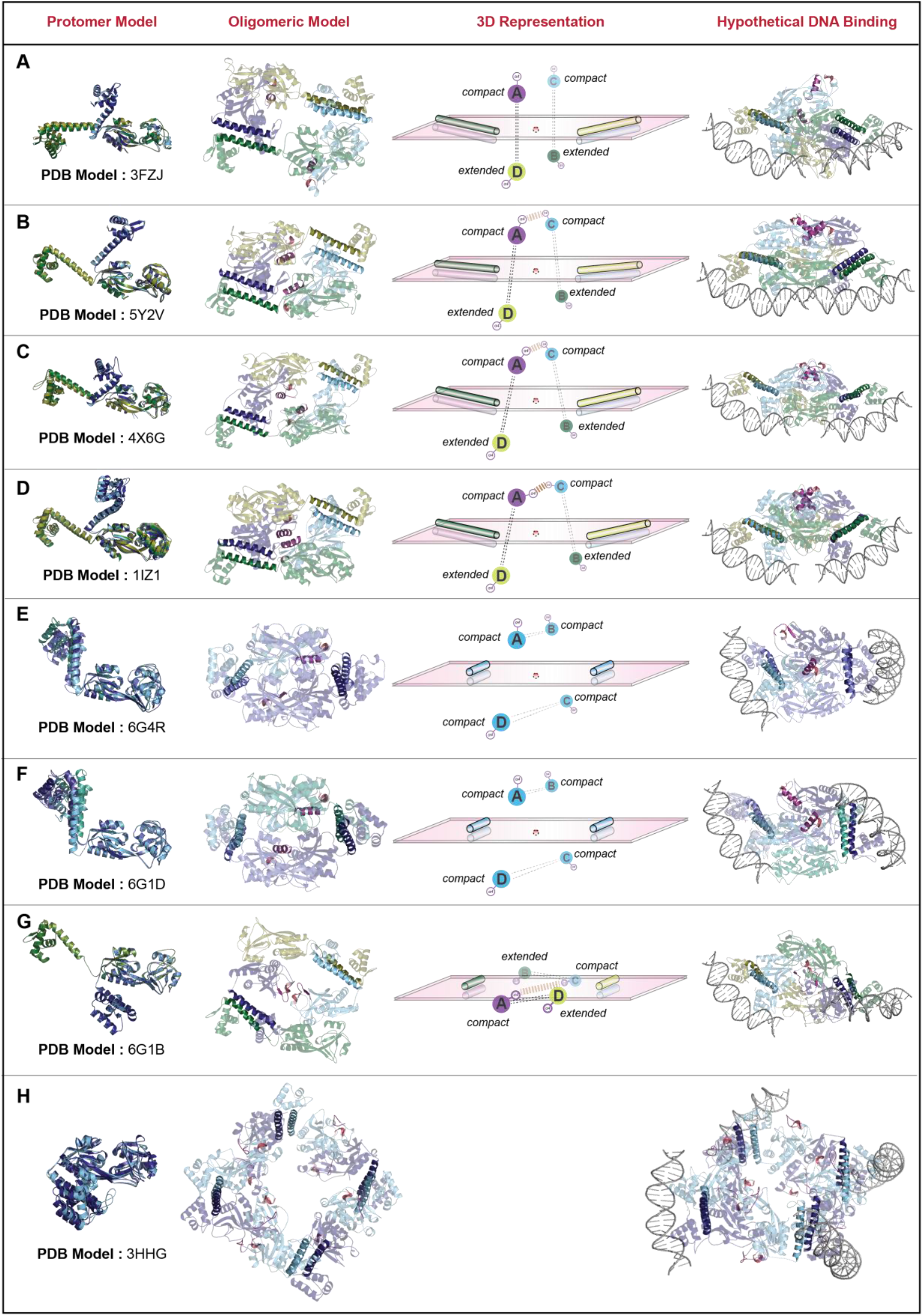
Model of Possible Oligomeric States of CynR, related to Figure 4. The CynR sequence (UniprotKB: P27111) was modelled from the indicated PDB structures that were used as a template, by the SWISS-MODEL server. The DNA in all structures was modelled from the structure of the HTH-DNA binding domain of CbnR in complex with the RBS of the cbnA promoter^3^. (**A-D**) Each tetramer is a dimer of dimers. The dimerization interface is formed by the long α-helix connecting the *CC* to the DNA binding domain (see also Fig. 4A, B). The α-helical coil-coil is stabilized always (i.e. in all known high-resolution structures of the HTH-motif transcription factors) by hydrophobic interactions. In one of the protomers, the helix is present in a compact configuration (i.e. it bends toward the *CC*), while in the other it is extended. The compact dimers are always represented by cold colors (blue-cyan), whereas the extended with warmer colors (green, yellow). The tetramerization interface is provided by two antiparallel *CCs* (Fig. S4B-E), one of them in the compact and the other in the extended configuration. By superimposing the *CC* of all protomers, we notice differences in the arrangement of the dimerization helix with respect to the *CC* between the compact or the extended protomers. This is clearly evident in the contact map (SuppFile. 7) of the protomers depicting the interactions between the dimerization helix and the *CC*, as such interactions vary even between the protomers having the same arrangement (either compact or extended). The dimerization helices are in the same plane, forming an intelligible rectangle, indicated in the 3D representation column. The two compact dimers are in one side on the rectangular plane, whereas the other two extended on the other (**3D Representation Column**). In some assemblies (4X6G and 6G1B), the quaternary structure is stabilized via interactions between secondary structure elements belonging to the compact dimers. Such elements (α4 and CH2, see SuppFile. 3) are colored with pink and magenta color respectively in all assemblies. The DNA is always present on the side of the two extended protomers. **(E, F)**: In these oligomers, all protomers are in the compact configuration, thus colored with cold colors. The DNA runs parallel to the intelligible rectangle denoting the plane of the dimerization helices. **(G)** Here two of the protomers are in the compact and two in the extended configuration alike A-D. The difference is that in the compact protomers the loop connecting the dimerization helix to the *CC* is rotated 180° with respect to the compact protomers of **A-D**, placing the DNA binding domain bellow the *CC*. In this case all *CCs* are in the same plane with that of the dimerization helices. **(H)** Here, all protomers are in the compact configuration, alike the compact protomers of panel G. With such an arrangement, an octameric assembly is formed. It has been verified that the octameric form is indeed the predominant form by native mass spectrometry and analytical ultracentrifugation.^4^

Strikingly, the quaternary assemblies presented in panels **E**, **F**, and **G** derive from the same protein at different conditions (*apo* vs. *liganded,* see also Discussion for details)^5^. Such structures indicate that the quaternary assembly that dictates DNA bending depends on the orientation of the dimerization helix with respect to the *CC*. No large domain motions of the D1-D2 of the protomers are observed is such quaternary assemblies (Table. S4). The differences between the protomers of these assemblies involve minor rearrangements of secondary structure elements and partial loss of secondary structure (in SuppFile. 3 denoted with a black rectangle). Moreover, such high-resolution data highlight the remarkable flexibility of the transcription factors, as the same protomer can obtain multiple conformations: two distinct compact configurations with the DNA binding domain either above or below the *CC* and even an extended configuration. There is even additional variability within the apparently similar compact and extended configurations (SuppFile. 7). Moreover, the prominent octameric population found for 6G1B^4^ (panel **G**) indicates that all of the protomers could also obtain the compact form of 3HHG (panel **H**). Only the compact form of 3HHG satisfies the geometrical criteria to obtain an octameric assembly.

This remarkable flexibility is also substantiated by inspecting the contact map of the CynR models based on the different oligomeric assemblies (SuppFile. 7). Such interaction map indicates that the dimerization helix could obtain multiple arrangements with respect to the *CC* of CynR in the different protomers, yielding multiple quaternary arrangements (**A-H**). This agrees with our HDX-MS results demonstrating that the dimerization helix and the DNA binding domain are extremely flexible (Fig. 4D).

The 3HHG octameric assembly is unique, as all other known LysR transcription factors are tetramers. Our phylogenetic tree indicated that this structure separates from all the rest in the first clade (SuppFile. 6). Moreover, the phylogenetic tree based uniquely on the *CCs* provides the same results. This means that the “information” for obtaining an octamer (i.e. all protomers in the compact orientation) resides within the *CC*. By inspecting the structure-based sequence alignment, we noticed that 3HHG has two differences compared to all other *CCPs* (SuppFile. 3): the additional helix CH5 connecting Hinge-β-sheet 2 (βH2) and CH6 and it lacks helix α4 that we represent with pink color and stabilizes tetrameric assemblies (see also Fig. 4B). The helix CH6 is placed just below the connecting-loop and fixes it in the compact configuration with the DNA binding domain below the *CC*. Helix α4 that stabilizes tetrameric assemblies is not required for formation of the octamer, and probably its absence facilitates octamerization as it destabilizes tetramerization.

**Figure S8:**
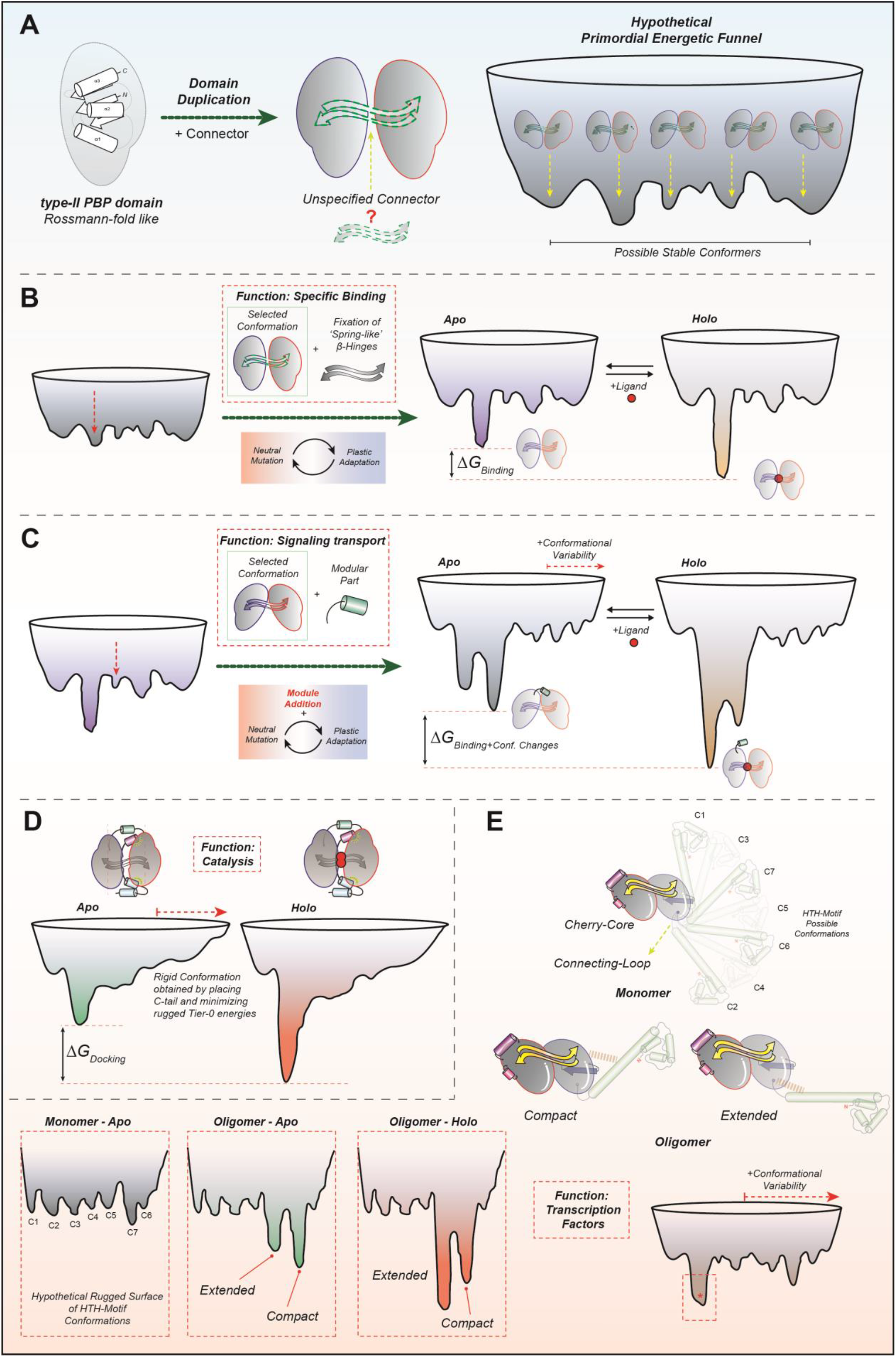
Hypothetical Energetic Funnels for the better visualization of the effects of Modularity on the Evolvability, related to Figure 6. **(A)** According to the view of generalists^6^, an ancestral structure composed of two interacting type-II PBP domains can acquire many distinct structural states depending on the arrangement of the two domains relative to each other. The energetic funnel of such an assembly will have many isoenergetic wells and manifest increased ruggedness, thus a high degree of ligand promiscuity (see also text for details). **(B)** Evolution of a generic ‘primordial’ two domain protein becoming a specialized *cherry-core*: When the two domains are linked with a β-sheet, a fix-length polypeptide chain emerges and a specific well (indicated with red arrow) is selected. However, it remains elusive if the evolution process is facilitated by utilizing either: (i) the positive selection (direct deepening of the well) or (ii) negative selection (destabilization of non-beneficial or unfunctional structures; indirectly deepen the well *relative* to other wells, becoming shallow). In the case of negative selection, the consecutive rounds of neutral mutations followed by plastic adaptations might indeed destabilized the wells of the non-functional structural states. The process results in the *relative* deepening and smoothening of the native state well (i.e. closed state, compared to the other wells) conferring ligand specificity (see discussion for details). The aforementioned structural arrangement represents the conserved *CC* structure. Solely the *CC* without embellishments is not encountered in extant proteins. Probably because the function of such a structure could be solely binding. As the *CC still* has the “remnants” of the highly-promiscuous ancestral two type-II PBP domains, we anticipate that this structure would exhibit a relatively high ligand promiscuity compared to the evolved *CCPs*. **(C-E)** Addition of the modular parts paves the way for evolution to sample a specific well on the funnel. If the presence of the module is beneficial for evolving the new function, nature may then introduce additional mutation to integrate the module into the core. Adopting the concept of protein evolvability, rounds of neutral mutations might be introduced synergistically on the module and the core and thus specify the well of the beneficial promiscuous structure. **(C)** A modular part represented by an asymmetric C-tail directs the evolutionary pressure to deepen one other specific well. Again, rounds of Neutral mutations coupled with plastic adaptations can greatly deepen this well, beyond the initial *CC* well (closed state), to create the open state well. This well is deep/smooth having increased ligand specificity. A distinct chemical environment represented by the presence of the ligand generates the *holo* energetic funnel. In the later one, the well corresponding to the closed state is the deeper. **(D)** A module consisting of a symmetric C-tail, directs evolution to deepen the closed state well and minimize the ruggedness of the funnel. This yields a very robust protein, providing the required stereochemistry necessary for specific reactions (see Fig. S4A). **(E)** Addition of an N-terminal domain having two modules (dimerization helix and DNA binding motif) widens the energetic funnel. In such a case a single closed well represents the *CC*, but additional ones derive from the flexible N-terminal domain. The *CC* is linked by a loop to the flexible domain, providing multiple wells in the energetic funnel (C1-C7). Oligomerization of such monomers will deepen specific wells in the oligomer funnel, similarly to the substrate in the *holo* funnel.

**Table. S1.** Large-scale Domain Motions (Tier-0 states) observed by comparing the *apo* and *holo* crystal structures. The domain movement comparison has been performed by the DynDom server using the indicated available crystal structures on the Protein Data Bank.

|  | ***Apo* Conformation**<br>**(PDB ID code)** | ***Holo* Conformation**<br>**(PDB ID code)** | **Moving Domain**<br>**of Core Scaffold** | **Rotation Angle**<br>**(deg)** | **Translation (Å)** |
| --- | --- | --- | --- | --- | --- |
| **Class A** | 2F7B | 2F7C | D2 | 4 | -0,4 |
|  | 3KOT | 4WKM | - | 0 | 0 |
|  | 4XWS | 4X6G | - | 0 | 0 |
|  | 2HXR | 3HFU | - | 0 | 0 |
| **Class B** | 4KR5 | 4KQP | D2 | 52,1 | 0,6 |
|  | 4P0I | 4POW | D2 | 47,7 | 1,8 |
|  | 3ZSF | 2YLN | D2 | 60,1 | -1,1 |
| **Class C** | 3L6G | 3L6H | D2 | 35 | -0,2 |
|  | 1SW5 | 1SW2 | D2 | 58,9 | -0,3 |
|  | 2REJ | 2RIN | - | 0 | 0 |
| **Class D** | 3S4U | 3QUJ | D2 | 74,3 | -0,5 |
| **Class E** | 2X7P | 2X7Q | D2 | 13,4 | -0,3 |
|  | 2I49 | 2I4B | - | 0 | 0 |
|  | 2DE2 | 2DE3 | - | 0 | 0 |
| **Class F** | 1SI1 | 1SI0 | D1 | 26,5 | -0,2 |
|  | 1O7T | 1D9Y | D1 | 19,7 | 0,1 |
|  | 1Y9U | 2OWT | D1 | 25,4 | -0,6 |
|  | 2HEU | 2I58 | D1 | 25,5 | 0,3 |
| **Class G** | 1OMP | 1ANF | D1 | 36 | -0,6 |
|  | 3OOA | 3OO6 | D1 | 37,9 | 0,3 |
|  | 5DVF | 5DVI | D1 | 18,9 | -0,2 |
|  | 1Y3Q | 1Y3N | D1 | 39,1 | 0,3 |
|  | 1J1N | 1KWH | D1 | 29,7 | 0,4 |

**Table. S2.**
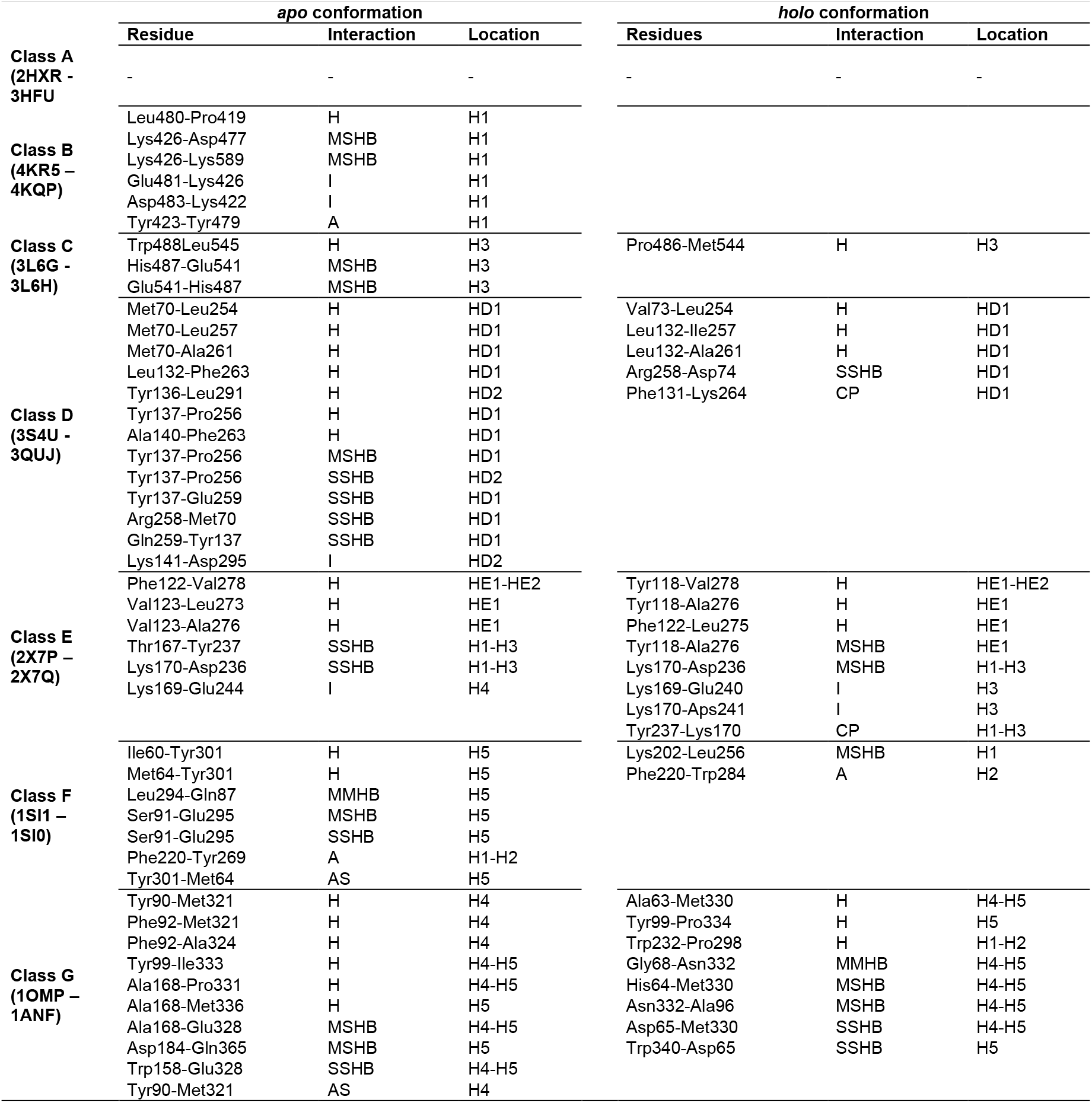
Stabilization Interactions of the Tier-0 *apo* and *holo* states. Only those interactions between the C-tail and D1/2 that specifically stabilize either the *apo* or *holo* state are shown. For each protein class the corresponding PDB ID code of the *apo* or *holo* structures is indicated. For class E, the only structures differing in the *apo* and *holo* states has been analyzed and shown. H (hydrophobic interaction, cut-off distance 5 Å), MSHB (main-chain side-chain hydrogen bond), I (ionic interaction, cut-off distance 6 Å), A (aromatic interaction, distance 4-7 Å), SSHB (side-chain side-chain hydrogen bond), MMHB (main-chain main-chain hydrogen bond), CP (cation-pi-interaction, cut-off distance 6 Å) and AS (aromatic-sulphur-interaction, cut-off distance 5 Å).

**Table. S3.** Apparent Melting Temperature Deduced from Intrinsic Trp Fluorescence of CynR. Intrinsic Trp fluorescence as a function of temperature (10°C-70°C) yielded a main T_m (apparent)_ and a secondary one characteristic of the presence of azide.

| | $T_m$ (°C) | | |
| --- | --- | --- | --- |
|  | Primary | Secondary | Azide |
| - DNA | 54.6 | N.A. | - |
|  | 55.3 | 26.9 | + |
| + DNA | 48.1 | N.A. | - |
|  | 48.3 | 26.5 | + |

**Table. S4.**
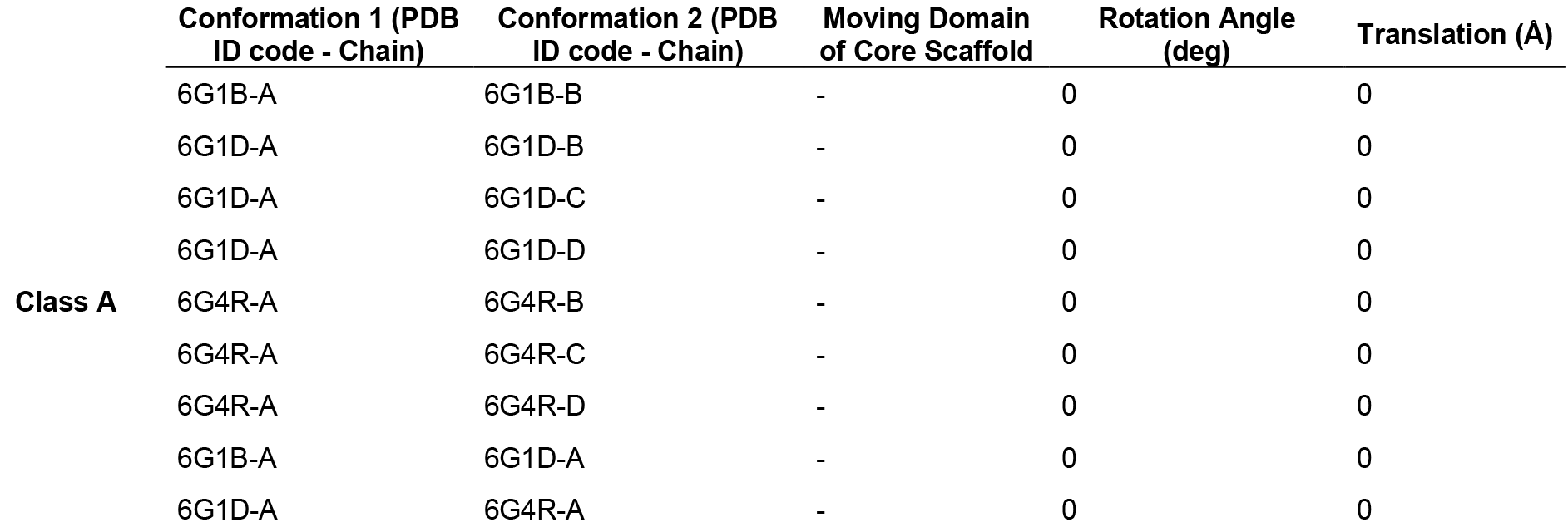
Tier-0 states of the indicated protomers within the tetrameric assemblies of the class A transcription factors. Analysis as described in Table. S1

**Table. S5.** Apparent FRET Efficiencies and Cα-Cα Distances derived from crystal structures. The apparent FRET efficiencies are the center and the errors are the limits of the 95% confidence interval of the Gaussian fit in **Fig. 3** and **5**. The PDB ID codes of proteins are reported in the Methods section and Table. S1.

|  | Apparent FRET efficiency |  | Ca- Ca distance (Å) |  |
| --- | --- | --- | --- | --- |
|  | Apo condition | Saturated condition | Open conformation | Closed conformation |
| CynR (C137 R233C) | 0.703 ± 0.003 | 0.707 ± 0.003 | 41.8 |  |
| CmpA (T89C N166C) | 0.439 ± 0.009 | 0.425 ± 0.010 | 48.2 |  |
| MalE (T36C S352C) | 0.586 ± 0.002 | 0.780 ± 0.002 | 51.3 | 40.8 |
| OpuAC (V360C V423C) | 0.550 ± 0.003 | 0.696 ± 0.003 | 45.7 | 36.3 |
| SBD2 (T369C S451C) | 0.586 ± 0.002 | 0.736 ± 0.002 | 49.0 | 40.1 |

## Legends of Tables Provided as Separate Files

**SFL1. Supplementary File S1 – Extended Sequence Based Phylogenetic tree**

Sequence-based extended phylogenetic tree based on percentage of sequence identity. Summary of this tree is presented in Fig. S1A. Every protein is represented by its PDB ID (known structures) or UniProt ID followed by its localisations identifier (SP, for presence of signal peptide; Lipidation, Glycosylation for lipidation and glycosylation signals respectively). If protein origin is not bacterial, origin (Archaea, Eukarya, Virus) is indicated.

**SFL2. Supplementary File S2 – Full Phylogenetic Trees**

Structure- and sequence-based phylogenetic trees of the 53 identified *cherry-core* proteins with known structures. Summary of this tree is presented in Fig. 1C and Fig. S1B-D. Trees are constructed after the entire polypeptide chain (**A**, **B**), the *CC* only (**C**) and the C-tail only (**D)**. Every protein is represented by its PDB ID code.

**SFL3. Supplementary File S3 – Secondary Structure Alignment Table**

<u>Sheet 1:</u> Revised alignments of *cherry-core* proteins. The identified proteins were sorted in different classes (**A-G**) depending on their N- & C-terminal regions (see also Fig. 1C, D). The alignment of the secondary structure elements is presented (**H**, for α-helix; **S**, for β-sheets) followed by the corresponding residues (the number of residues forming the secondary structural element is indicated in parenthesis). Every protein is represented by its PDB ID code followed by its chain identifier. We represent equivalent secondary structure elements (occupying the same place in the 3D space) with the same uniform color throughout a column. We identified distinct protein classes **A-G** according to the C-terminal regions of their CC, which is formed by different secondary structure elements. Secondary structure elements that are joined (i.e. two α-helices becoming a long one) are marked by red letters. We indicated subclasses with asterisks. Subclasses represent proteins having minor differences in the secondary structure elements of their C-tails, with respect to all other proteins of the same class, e.g., A* has an additional terminal helix at the C-tail similar to H1, but positioned in a different location in the 3D space compared to A. Similarly, for B* which has an additional helix (CH5) and an extra small β-sheet compared to B. The class B** has two extremely short helices CH6 and H1, while sub-class B*** and G* have one additional helix at the C-tail with respect to other B and G class members, respectively.

<u>Sheet 2:</u> Classification of the proteins included in Sheet 1 according to the ECOD database.

**SFL4. Supplementary File S4 – HDX MS Data for CynR**

Excel file composed of 4 sheets containing 3 tables (F4A, F4B, F4C) and coverage maps. The sheets include detailed description of the conditions and analysis of the HDX-MS experiments.

**SFL5. Supplementary File S5 – HDX MS Data for MalE**

Excel file composed of 7 sheets containing 5 tables (F5A, F5B, F5C, F5D, F5E), difference plots and coverage maps. The sheets include detailed description of the conditions and analysis of the HDX-MS experiments.

**SFL6. Supplementary File S7– Structure based phylogenetic trees of Class A Protein**

Results as obtained from the Dali server (see Methods). The structure based phylogenetic tree of class A proteins with known oligomeric structures (see Fig. S7). Top part contains full length class A transcription factors (HTH motif/dimerization helix and CC) while bottom part, only the *CC*.

**SFL7. Supplementary File S7–Interaction of CynR connecting loop/dimerization helix with the CC.**

The *cynR* sequence was modelled after the known oligomeric high-resolution structures (see Fig. S7 and methods section). The contact interfaces between the dimerization helix and the *CC* derived from the structures (left part of the excel table) or from the corresponding models (right part of the excel table) were analyzed in the Protein Interaction calculator server, using standard server settings (alike Table S2, see also Methods). The residues forming such interface belong to the connecting-loop or the dimerization helix (indicated as loop and helix respectively) and elements of the *CC* (indicated according to the SuppFile. 3). The nature of the interactions, hydrophobic, aromatic, Cation-P_i_, hydrogen-bonds, Ionic are represented with blue, cyan, green, orange and red color respectively.

